# The molecular principles of Piwi-mediated co-transcriptional silencing through the dimeric SFiNX complex

**DOI:** 10.1101/2021.01.08.425619

**Authors:** Jakob Schnabl, Juncheng Wang, Ulrich Hohmann, Maja Gehre, Julia Batki, Veselin I. Andreev, Kim Purkhauser, Nina Fasching, Peter Duchek, Maria Novatchkova, Karl Mechtler, Clemens Plaschka, Dinshaw J. Patel, Julius Brennecke

## Abstract

Nuclear Argonaute proteins, guided to nascent target RNAs by their bound small RNAs, elicit co-transcriptional silencing through heterochromatin formation at transposon insertions and repetitive genomic loci. The molecular mechanisms involved in this process are incompletely understood. Here, we propose that the SFiNX complex, a silencing mediator downstream of nuclear Piwi-piRNA complexes in *Drosophila*, enables co-transcriptional silencing via the formation of molecular condensates. Condensate formation is stimulated by nucleic acid binding and requires SFiNX dimerization, mediated by the dynein light chain protein, LC8/Cutup. LC8’s function within SFiNX can be bypassed with a heterologous dimerization domain, suggesting that dimerization is a constitutive feature of SFiNX. Mutations preventing LC8-mediated SFiNX dimerization result in loss of condensate formation *in vitro* and inability of Piwi to initiate heterochromatin formation and silence transposons *in vivo*. Formation of molecular condensates might be a general mechanism that underlies effective heterochromatin establishment at small RNA target loci in a co-transcriptional manner.

## Introduction

Heterochromatin is critical for maintaining genome integrity in eukaryotes, suppressing the transcription of repetitive sequences and preventing them from undergoing ectopic recombination^1-4^. Initiation and maintenance of heterochromatin require post-translational modifications of histone tails, in particular the deposition of repressive di- and trimethyl groups on Lysine 9 of histone H3 (H3K9me2/3)^5,6^. As a consequence, various proteins with an affinity for H3K9me2/3 are recruited and stabilized on chromatin^7-9^. Their biophysical and/or enzymatic activities lead to additional histone modifications, reduced nucleosome dynamics and increased chromatin compaction^5^.

Heterochromatin plays a central role in the transcriptional silencing of transposons, selfish genetic elements that make up large parts of plant, fungal, and animal genomes^2-4^. Transposon insertions include features that resemble endogenous gene loci, requiring mechanisms that direct the heterochromatin machinery specifically to these sites. Two strategies that enable sequence-specific heterochromatin formation are conserved in eukaryotes^10^. The first is based on DNA-binding repressor proteins, which recognize specific motifs in transposon sequences^11^, and is independent of target locus transcription. The second strategy depends on target locus transcription and is referred to as co-transcriptional gene silencing. It involves small RNAs binding to nuclear Argonaute proteins^12^ and guiding them to complementary, nascent transposon transcripts in chromatin^13-16^.

In most animals, co-transcriptional silencing of transposable elements is associated with PIWI-clade Argonaute proteins, which are complexed with PIWI-interacting RNAs (piRNAs)^17-19^. In the *Drosophila melanogaster* ovary, piRNAs encoded by genomic loci that provide heritable libraries of transposon sequences are loaded into three PIWI-clade Argonautes^20-22^. One of them, Piwi, acts in the nucleus^23-25^. Piwi-piRNA complexes are recruited to complementary, nascent transposon transcripts and orchestrate cotranscriptional heterochromatin formation at hundreds of transposon insertions genome-wide^26-29^.

Piwi-mediated silencing requires several general heterochromatin factors. Among those are H3K9 methyltransferases, histone deacetylases, chromatin remodelers, the SUMOylation machinery and HP1 proteins that bind the H3K9me2/3 mark and mediate target locus compaction^30-34^. According to current models, only target-engaged Piwi–piRNA complexes are capable of recruiting heterochromatin effectors via dedicated bridging factors^30,31^. A central bridging factor in this context is the SFiNX complex (a.k.a. Pandas, PICTS or PNNP complex)^35-38^. SFiNX is required for piRNA-guided co-transcriptional silencing, and its experimental tethering to nascent RNA is sufficient to induce heterochromatin formation, independent of Piwi and piRNAs^30,31^. The SFiNX complex consists of the orphan protein Panoramix (Panx) and the heterodimeric nuclear RNA export variant Nxf2–Nxt1^30,31,35-38^. SFiNX has two separable activities. The first involves signaling to the silencing effectors and resides within the N-terminal, low-complexity region of Panx^37^. Experimental recruitment of this ∼200 amino acid peptide directly to a DNA locus is sufficient to induce strong silencing via unknown mechanisms. The same peptide, however, is unable to elicit silencing when recruited to chromatin via a nascent target RNA. Co-transcriptional, RNA-directed silencing requires the second SFiNX activity. It resides in the structured, C-terminal half of Panx that interacts with the Nxf2– Nxt1 heterodimer^35^, and its molecular nature is unknown. These observations suggest there are fundamental differences in heterochromatin formation downstream of DNA-binding versus RNA-binding factors. Indeed, chromatin regulators such as the H3K4 demethylase Lsd1, the H3K9 methyl-transferase SetDB1, or the H3K9me2/3 binding protein HP1 are potent silencers when recruited to a reporter locus via DNA-targeting but not via targeting to the nascent RNA^30,31,35-38^.

Here, we reveal that SFiNX forms molecular condensates *in vitro* in a process that is enhanced by nucleic acid binding. Mutations that impair condensate formation abrogate SFiNX’s co-transcriptional silencing capacity *in vivo*. Nucleic acid binding and condensate formation depend strictly on homo-dimerization of SFiNX, which is mediated by dynein light chain 8 (LC8), a highly conserved dimerization hub protein that we identify as fourth integral SFiNX subunit. We propose that SFiNX-mediated formation of molecular condensates enables piRNA-guided co-transcriptional silencing, possibly by retaining the target RNA and thus the recruited silencing complexes at chromatin.

## Results

### The dynein light chain LC8 is a SFiNX complex subunit

With the aim to study SFiNX biochemically, we set out to determine all of its core subunits. We immuno-purified SFiNX from the nuclear lysate of cultured ovarian somatic cells (OSCs)^39,40^ expressing the SFiNX subunit Panx from its endogenous locus and fused to a GFP-FLAG tag (Figure 1A). Silver staining of the immuno-precipitate revealed four prominent protein bands. Three corresponded to the known SFiNX subunits Panx, Nxf2 and Nxt1. The fourth corresponded to Cutup, the 8 kDa Dynein Light Chain 8 (LC8) ortholog of *Drosophila*^41,42^, suggesting that LC8 is a SFiNX subunit.

**Fig. 1.**
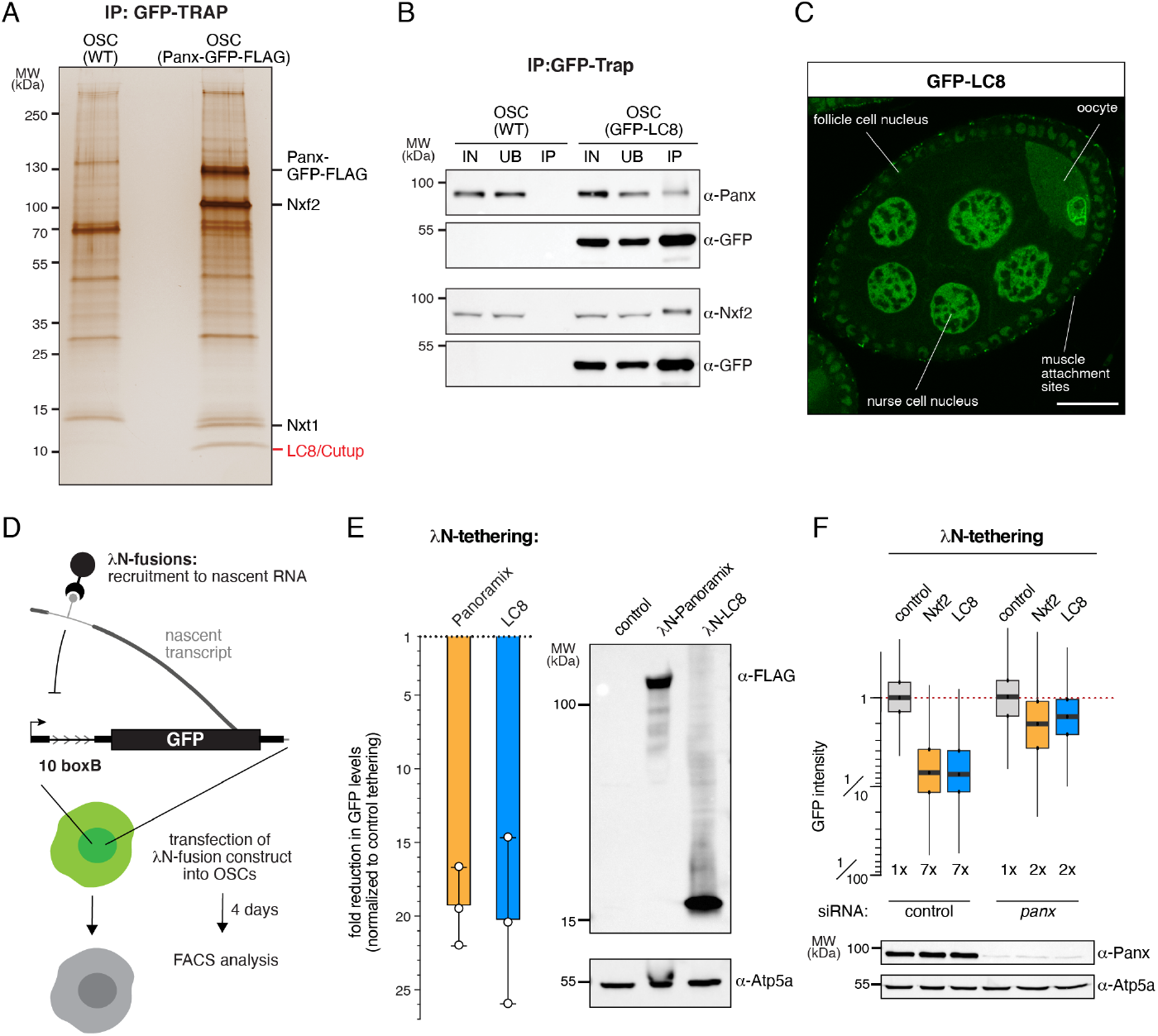
The dynein light chain Cutup/LC8 is an integral SFiNX complex subunit. **A**, Silver stained SDS-PAGE (molecular weight marker to the left) of immunoprecipitated Panx (right lane; from endogenously FLAG-GFP-tagged cell line) and control (left lane; from wildtype OSCs). **B**, Western blot analysis indicating co-immuno-precipitation of endogenously tagged LC8 from OSCs with endogenous Panx and Nxf2 (wildtype OSCs served as control). **C**, Confocal image of *Drosophila* egg chamber showing GFP-tagged LC8 (scale bar: 20µm). **D**, Cartoon showing the co-transcriptional silencing reporter system in OSCs. **E**, Bar diagram (left) indicates fold repression of the GFP-reporter after transfection of indicated λN-tagged proteins normalized to control (data represent mean +/-St.dev. of 3 biol. replicates). Western blot analysis (right) showing expression of indicated λN fusion proteins in OSCs (Atp5a served as loading control). **F**, Box plots showing GFP reporter intensity in OSCs 5 days after transfection with siRNAs against control or *panx* and 3 days after transfection with plasmids expressing indicated λN fusion proteins (top, numbers indicate fold-repression values normalized to control; box plots indicate median, first and third quartiles (box), whiskers show 1.5× interquartile range; outliers were omitted; n=2500 cells). The western blot below shows Panx protein levels in control versus *panx* siRNA transfected OSCs (Atp5a served as loading control).

LC8 was first described as a component of the cytoplasmic dynein multi-protein complex, a minus-end directed microtubule motor^43,44^. SFiNX instead is a nuclear complex. To confirm the LC8–SFiNX interaction we carried out reciprocal co-immuno-precipitation experiments using lysate from OSCs expressing endogenously FLAG-GFP tagged LC8 (Figure S1A). The SFiNX subunits Panx and Nxf2 robustly and specifically co-purified with tagged LC8 (Figure 1B). To determine the cellular distribution of LC8 *in vivo*, we analyzed a fly line expressing GFP-tagged LC8 from a genomic transgene. In ovaries, LC8 was enriched in foci at the basal surface of the follicular epithelium (potentially muscle attachment sites) and in cytoplasmic foci of unknown origin in germline nurse cells and the oocyte (Figure 1C). A substantial fraction of GFP-LC8, however, localized to the nucleus in ovarian somatic and germline cells. A similar pattern was seen in other tissues such as wing imaginal discs or the larval fat body (Figure S1B). These findings were consistent with a physical LC8-SFiNX interaction and pointed to a general role of LC8 in nuclear processes^45^.

To investigate whether the interaction between LC8 and Panx– Nxf2–Nxt1 implicates LC8 in the piRNA pathway, we used a cotranscriptional silencing assay in OSCs where the silencing capacity of a protein of interest can be determined by recruiting it via the λN-boxB system to nascent transcripts expressed from a stably integrated GFP-reporter transgene (Figure 1D)^30,31^. Tethering of the SFiNX subunits Panx or Nxf2 results in ∼20-fold reporter silencing via heterochromatin formation^35^. Recruitment of LC8 led to a similarly strong repression (Figure 1E). Reporter silencing by LC8 was dependent on SFiNX as depletion of Panx abrogated the silencing response elicited by λN-LC8 recruitment, as also seen for tethering of λN-Nxf2 (Figure 1F; S1C). Thus, LC8 is a fourth core subunit of the SFiNX complex.

### LC8 is required for SFiNX function

Like for other piRNA pathway genes, *panx* mutant flies are viable but sterile, exhibiting strong transposon de-repression. In contrast, mutations in the ubiquitously expressed *LC8* gene are lethal^41,42^. To ask whether LC8 is required for piRNA pathway function, we depleted LC8 in OSCs using RNAi. We observed the de-repression of *gypsy* and *mdg1*, two LTR retroelements that are under piRNA pathway control in OSCs (Figure 2A, B). In addition, Panx and Nxf2 protein levels (but not mRNA levels) were reduced in LC8 depleted versus wildtype cells (Figure 2B; S2A). Considering the reciprocal stabilization of Panx and Nxf2^35-38^ our data suggest that LC8 is required for the function as well as for the assembly of the SFiNX complex.

**Fig. 2.**
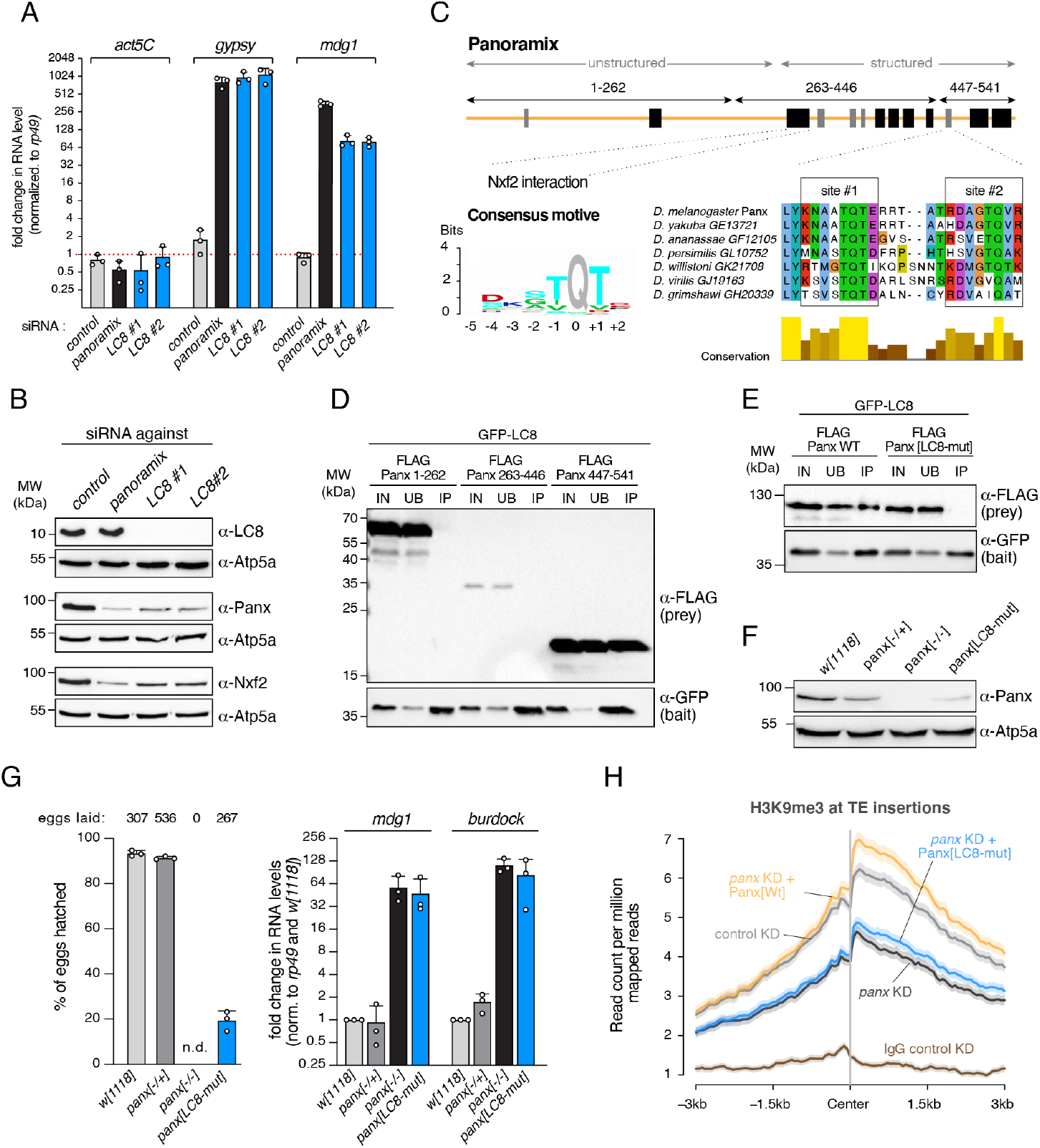
LC8 is required for SFiNX-mediated transposon silencing. **A**, Bar graph showing mRNA levels (normalized to *rp49* levels) of the endogenous LTR retro-transposons *gypsy* and *mdg1* after depletion of Panx or LC8 (two independent siRNAs). *actin5C* mRNA levels served as control. Data represent mean +/-St.dev. of 3 independent experiments. **B**, Western blot showing protein levels of indicated proteins after depletion of Panx or LC8 (two independent siRNAs; Atp5a served as loading control). **C**, Schematic representation of Panx, indicating predicted secondary structure elements (black: alpha-helices, grey: beta-strands) and the two LC8 binding sites and their evolutionary conservation below. The sequence logo to the left depicts the consensus LC8 binding motif as determined in [73]. Protein regions tested in D, are indicated. **D, E**, Western blot analyses of GFP-LC8 immuno-precipitation experiments using lysate from S2 cells transiently transfected with indicated FLAG-Panoramix expressing plasmids (relative amount loaded in immuno-precipitation lanes: 4x; IN: input, UB: unbound, IP; immuno-precipitate). **F**, Western blot showing Panx protein levels in ovarian lysates from flies of indicated genotype (Atp5a served as loading control). **G**, The left bar graph shows hatching rates of eggs laid by flies of indicated genotype (data represent mean + St.dev. of 3 independent replicates. The right bar graph shows mRNA levels of somatic (*mdg1*) and germline (*burdock*) transposons in total ovarian RNA from flies of indicated genotype (normalized to *rp49* and control flies; data represent mean + St.dev. of 3 independent replicates). **H**, Meta profiles showing H3K9me3 levels (determined by Cut & Run) in a 6kb window around 381 piRNA-controlled transposon insertions (center) in OSCs transfected with indicated siRNA (control or *panx* KD) and rescue constructs (IgG served as control).

LC8 has been implicated in diverse dynein-dependent and independent cellular processes^46,47^. The transposon silencing defects in LC8-depleted OSCs might therefore be indirect. To determine whether LC8 is directly required for SFiNX function, we first defined the physical interaction between LC8 and the other SFiNX subunits. LC8 proteins form stable homodimers that interact with many cellular proteins. The dimer interface generates two symmetric binding grooves for peptides, which harbor a consensus motif with a ‘TQT’ anchor and which adopt a beta-strand conformation upon binding to the LC8 dimer^46,47^ (Figure 2C). To determine whether Panx, Nxf2 or Nxt1 interact directly with LC8, we used established computational tools^45,48^ to predict potential LC8-interacting peptides in each of the three other subunits. Two such binding motifs were identified within a short, disordered stretch of the C-terminal portion of Panx (NAA**<u>TQT</u>**E and DAG**<u>TQV</u>**R) and both sites are evolutionarily conserved (Figure 2C). To test whether Panx interacts with LC8 via the predicted binding motifs, we performed co-IP experiments using lysates from *Drosophila* Schneider 2 cells (S2) transiently expressing GFP-LC8 and FLAG-Panx. Only the C-terminal portion of Panx (AA447-541), including the two predicted binding sites, interacted with LC8 (Figure 2D). As this region does not harbor the binding site for the Nxf2–Nxt1 heterodimer, these results establish Panx as the LC8 interaction partner in SFiNX. In order to disrupt the LC8–Panx interaction, we introduced alanine substitutions into the TQT core motifs in Panx. Mutation of both sites, but not of either site alone, resulted in loss of the interaction (Figure 2E, S2B). Thus, the two motifs in Panx are independent LC8 binding sites.

To determine whether LC8 is directly required for Panx and, therefore, SFiNX function we used CRISPR/Cas9 to generate flies harboring alanine substitutions in the TQT core of both LC8 interaction sites in the endogenous *panx* locus (*panx[LC8-mut]*) (Figure S2C). Homozygous *panx[LC8-mut]* flies were viable but exhibited reduced Panx protein levels (Figure 2F), consistent with the reduced Panx levels seen in OSCs lacking LC8 (Figure 2B). *panx[LC8-mut]* flies also showed reduced fertility (egg hatching rate: ∼ 20%) and displayed transposon de-repression in the ovarian soma and germline to an extent resembling that of *panx* null mutants (Figure 2G). In comparison, heterozygous *panx[+/-]* flies, which have Panx protein levels comparable to the *panx[LC8-mut]* mutant, exhibited no defects in transposon silencing and fertility (Figure 2F, G). Consistent with the strong transposon silencing defects, OSCs expressing Panx[LC8-mut] instead of the wildtype protein displayed reduced levels of H3K9me3 around piRNA-controlled transposon insertions (Figure 2H)^27,31^. Taken together, our data establish that LC8 is directly required for SFiNX function.

### LC8 does not connect SFiNX to other protein complexes

LC8 has originally been proposed to function as cargo adaptor for the dynein motor complex^49,50^. We hypothesized that LC8 might analogously recruit downstream silencing effectors of the piRNA pathway to SFiNX. To identify LC8 interaction partners, we performed co-IP experiments with FLAG-GFP tagged LC8 from OSC whole cell lysates including an endogenously modified *LC8* locus and from fly ovaries expressing tagged LC8 from a genomic transgene (Figure 1C, S1A). Quantitative, label-free mass spectrometry revealed a large number of proteins co-purifying with LC8 (Figure 3A, B). We identified 19 of 30 annotated LC8 interaction partners in Flybase^51^, including proteins linked to the dynein motor complex (Sw, Dlc90F, Robl), the microtubule linked protein Kank, and the centriolar protein Ana2 (Table S1). In addition to known cytoplasmic interactors, several nuclear proteins were enriched in LC8 immuno-precipitates. Besides SFiNX subunits Panx, Nxf2, and Nxt1, these included a complex involved in telomer biology and transcriptional regulation (Woc, Row, HP1b, HP1c)^52,53^, the Tousled-like kinase (regulator of replication-dependent chromatin assembly) ^54^, and the H3K9 methyltransferase complex consisting of Eggless/SetDB1 and its partner proteins Windei/ATF7IP and CG14464/ARLE-14^55-57^, suggesting an important role for LC8 in chromatin-related processes.

**Fig. 3.**
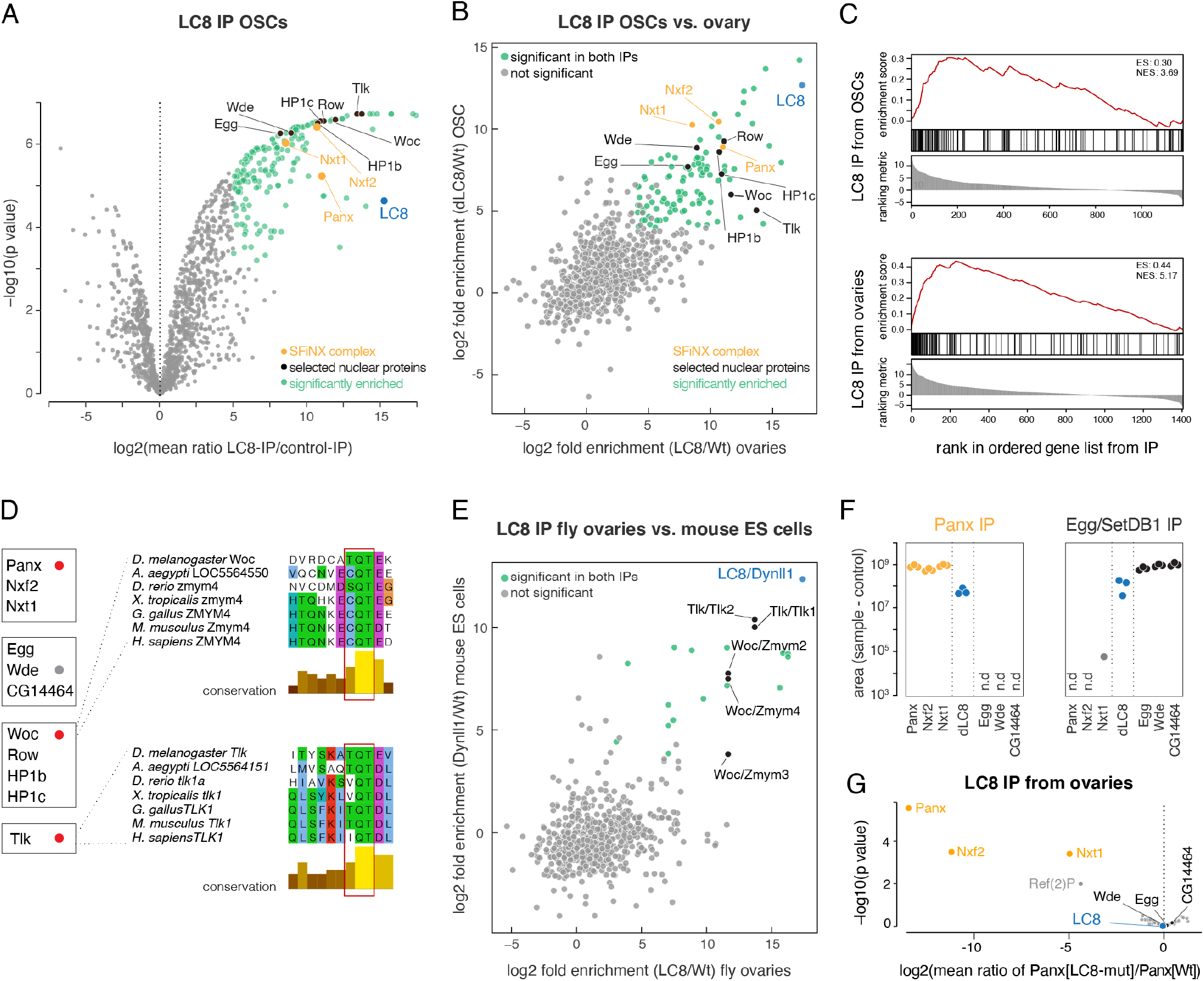
LC8 mediates dimerization of diverse nuclear protein complexes. **A**, Volcano plot showing fold enrichments versus statistical significance (determined by quantitative mass spectrometry) of proteins in FLAG-GFP LC8 co-immuno-precipitates versus control (n=3 biological replicates; experimental OSCs express endogenously tagged LC8; wildtype OSCs served as control). The bait (LC8) as well as selected interacting nuclear proteins are labelled. **B**, Scatter plot showing fold enrichments (versus control) of proteins co-immuno-precipitating with LC8 from OSC versus ovary whole cell lysates. The bait (LC8) as well as selected interacting nuclear proteins are labelled. **C**, Enrichment of predicted LC8 motives in LC8 co-purifying proteins (top: from OSCs; bottom: from ovaries) determined by gene set enrichment analysis. Protein lists were ranked by their fold enrichment (ranking metric) in the respective co-immuno-precipitation experiments. LC8-hub45 was used for LC8 motive prediction. **D**, Selected nuclear LC8 interactors grouped into known protein complexes (dots indicate predicted LC8 interaction motives; red: experimentally confirmed; gray: experimentally not confirmed). To the right, protein sequence alignments of indicated LC8 binding sites are shown; TQT motives (mutated in Fig S3A, FigS3B) are highlighted with red boxes. **E**, Scatter plot showing fold enrichments (versus control) of proteins co-immunoprecipitating with LC8 (from OSCs) or with Dynll1 from mouse ES cells. The bait (LC8/Dynll1) as well as selected interacting nuclear proteins are labelled. **F**, Absolute peak intensities of peptides from indicated proteins in Panx or Egg co-immuno-precipitates (values of matched controls were subtracted from experimental values; n=3 replicates; n.d.: not detected). **G**, Volcano plot showing fold enrichment of proteins (determined by quantitative mass spectrometry) in FLAG-GFP-LC8 co-immuno-precipitates from *panx[LC8-mut]* versus wildtype ovaries. (n=3 biological replicates).

To understand how LC8 binds such a diverse set of proteins we explored whether the identified interactors contained ‘TQT’ binding motifs. LC8 co-purifying factors were strongly enriched in predicted high-scoring LC8 binding motifs (Figure 3C). For most of the protein complexes analyzed, only one subunit harbored a predicted LC8 binding motif. For SFiNX, this was Panx. For the Woc–Row– HP1b–HP1c complex, this was Woc, the fly ortholog of the mammalian Zmym2/3/4 transcriptional regulators, and mutation of the TQT motif disrupted the physical interaction with LC8 (Figure S3A). Similar results were obtained for Tousled-like kinase (Figure S3B). Our data are consistent with findings in other species indicating that the LC8 homodimer acts as a binding partner for various client proteins possessing an accessible TQT binding motif^46,47^.

LC8 is highly conserved, with the fly and human proteins being 96% identical over their entire length. To investigate whether LC8’s interaction with nuclear factors is also conserved, we performed co-IP experiments in mouse ES (mES) cells. Mouse LC8/Dynll1 is present in both cytoplasm and nucleus, and LC8/Dynll1 co-immuno-precipitates with Zmym2/4 (the mouse ortholog of Woc) and the Tousled-like Kinases (Figure 3E; S3C, D). The LC8 binding sites in Woc/Zmym2/4 and Tousled-like kinase are evolutionarily conserved from flies to mammals (Figure 3D), indicating that LC8’s function in the nucleus is ancient and not connected solely to SFiNX, a protein complex specific to diptera.

LC8’s ability to bind diverse nuclear protein complexes might function to connect SFiNX to defined piRNA-pathway downstream chromatin effectors. We noted that the Eggless/SetDB1 complex, a H3K9 methyltransferase involved in heterochromatin establishment, was highly enriched in the LC8 co-immuno-precipitation (Figure 3A, B). SetDB1 and its cofactor Windei/Atf7IP act downstream of SFiNX during piRNA-guided transposon silencing and heterochromatin formation in OSCs and fly ovaries^30,31,55^. To assay for a physical interaction between SFiNX and the SetDB1 complex (potentially mediated by LC8) we used quantitative mass spectrometry to identify proteins co-purifying with SFiNX (using endogenously tagged Panx) or the SetDB1 complex (using endogenously tagged Eggless; Figure S3E). LC8 was among the highest co-enriched proteins in both experiments. However, SFiNX subunits were not present in the SetDB1 complex purification and vice versa (Figure 3F, S3F-G). This strongly suggests that LC8 binds independently to the two protein complexes.

To further test this prediction, we performed LC8 co-immunoprecipitation experiments from ovary lysates of wildtype flies versus flies expressing *panx[LC8-mut]*. Among the 115 proteins that were highly enriched (>30-fold) in LC8 immuno-precipitates (Figure 3B), only SFiNX subunits Panx, Nxf2, and Nxt1 failed to interact with LC8 in *panx[LC8-mut]* lysates while LC8 binding to the SetDB1 complex was unchanged (Figure 3G). Altogether, our findings do not support a model where LC8 connects SFiNX to down-stream heterochromatin effectors. Instead, they suggest that LC8 is part of several, physically separate cytoplasmic and nuclear protein complexes. Considering that the LC8 homodimer harbors two symmetric binding sites for TQT motif-containing peptides, our data support a role for LC8 as a versatile homo-dimerization module for various cellular protein complexes^45-47^.

### The SFiNX complex is a homodimer

To test whether SFiNX forms LC8-mediated dimers or oligomers, we used the OSC line harboring a heterozygous, tagged *panx* allele and performed co-immuno-precipitation followed by western blot. This revealed that tagged and un-tagged Panx co-purify (Figure 4A). Similarly, transiently expressed GFP-Panx and FLAG-Panx interact, and this interaction was dependent on the two LC8 binding sites in Panx, indicating that LC8 is required for SFiNX oligomerization (Figure S4A).

**Fig. 4.**
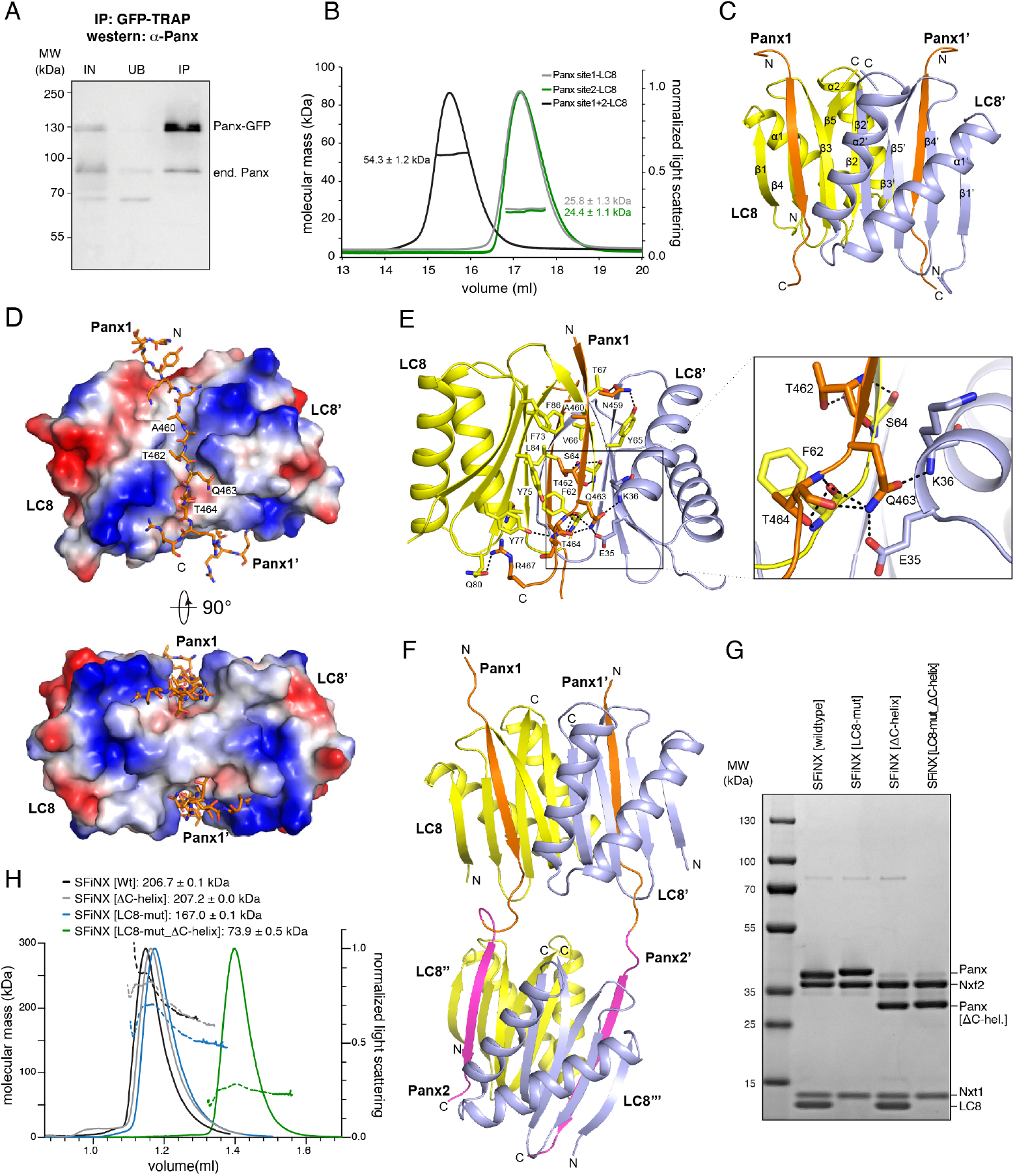
The SFiNX complex is a homodimer. **A**, Western blot analysis of a Panx-GFP immunoprecipitation experiment from OSCs harboring one GFP-tagged and one wildtype *panx* allele (IN: input, UB: unbound, IP; immuno-precipitate). **B**, Molecular weight determination of LC8 in complex with Panoramix peptides containing LC8 binding site#1 (TQT site), site#2 (TQV site) or both sites as measured by SEC-MALS. **C**, Ribbon representation of the structure of a LC8 homodimer (monomers in yellow and cyan) in complex with two Panoramix peptides containing the LC8 interacting site#1 (TQT site). **D**, Electrostatic surface representations of the LC8 homodimer in complex with two Panoramix peptides containing the LC8 interacting site#1 (TQT site; shown in stick representation). **E**, Intermolecular interactions (hydro-phobic and hydrogen bonding networks) between the LC8 dimer (monomers in yellow and cyan) and a Panoramix peptide containing the LC8 interacting site#1 (orange). The TQT motif recognition is shown in the closeup. **F**, Ribbon representation of the structure of two LC8 dimers (yellow and blue) in complex with two Panoramix peptides containing LC8 interacting sites #1 and #2. **G**, Coomassie stained SDS PAGE showing recombinant SFiNX complex variants with indicated protein subunits analyzed in H. **H**, SEC-MALS chromatogram and determined molecular weight of indicated recombinant SFiNX complex variants (from G).

To more precisely characterize the oligomerization mode of Panx, we purified LC8 together with Panx peptides harboring either TQT site #1, site #2, or both. Size exclusion chromatography coupled to multi-angle light scattering (SEC-MALS) revealed that the molecular weight of the protein complexes corresponded to either one LC8 homodimer with two single-TQT Panx peptides (predicted MW: 23.5 and 23.8 kDa) or two LC8 homodimers with two dual-TQT Panx peptides (predicted MW: 47.5 kDa) (Figure 4B). Isothermal titration calorimetry (ITC) measurements revealed comparable affinities for the two individual TQT sites in Panx (TQT-peptide: KD ∼ 75nM; TQV-peptide: KD ∼ 140nM), in line with previous measurements for unrelated LC8-peptide complexes^58^. A Panx peptide with both core TQT motifs mutated to alanine showed no binding to LC8 (Figure S4B). Thus, LC8 mediates homo-dimerization of Panx (and SFiNX) via two independent TQT motifs and does not promote the formation of higher order oligomers, even though this is theoretically possible given the two tandem LC8 binding sites in Panx.

Using the recombinant LC8–Panx[single TQT-site] complexes, we determined their atomic structures at 1.42 Å resolution for site #1 and 1.79 Å resolution for site #2, respectively (Figure 4C, S4C; Table S2). TQT peptides #1 and #2 adopt an almost identical binding mode to LC8 that is similar to known LC8–peptide complexes^59,60^: the LC8 dimer contains two hydrophobic-lined grooves on opposite faces, allowing two parallel Panx peptides to bind (Fig. 4D). The TQT motif is recognized through a hydrogen-bonding network involving residues from both LC8 monomers (Fig. 4E). To determine the conformation of LC8 with tandem TQT sites, we solved the crystal structure of the LC8–Panx [dual TQT-site] complex at 2.5 Å resolution (Fig. 4F, Table S2). This structure showed that two LC8 homodimers bind the two closely spaced TQT motifs from a pair of parallel aligned Panx peptides. The LC8–Panx interaction for each site is highly similar to the complexes harboring single TQT peptides (with RMSDs of 0.25 for peptide #1 and 0.31 for peptide #2). Our structural analyses confirm that the two tandem TQT motifs in Panx are compatible with a homo-dimerization via two independent LC8 dimers.

To test if LC8-mediated dimerization of Panx also occurs in the context of the SFiNX complex, we purified recombinant SFiNX consisting of Panx[263-541], Nxf2[541-841], Nxt1, and LC8 from insect cells. This complex lacks the N-terminal low-complexity region of Panx (implicated in silencing) and the N-terminal double RRM-LRR domains in Nxf2 (implicated in RNA binding)^35-38^. All four proteins formed a stable, uniform complex with a molecular weight corresponding to a homodimer consisting of two molecules each of Panx, Nxf2 and Nxt1 plus four molecules of LC8. (Figure 4G, H; note the ∼ twofold intensity of the LC8 band compared to the Nxt1 band in the Coomassie stained SDS gel).

We previously reported that the ternary protein complex consisting of Nxf2–Nxt1 and C-terminally truncated Panx is monomeric^35^, which is to be expected as, in this complex, Panx lacks the C-terminal 95 amino acids, including both TQT motifs. We hypothesized that SFiNX should also be monomeric when co-expression of LC8 is omitted, even when the Panx C-terminus is intact. As expression of wildtype Panx led to the co-purification of endogenous LC8 from insect cells, we expressed the Panx[LC8-mut] variant for this experiment. The ternary Panx[263-541 LC8-mut]–Nxf2[541-841]–Nxt1 complex, despite lacking LC8, was still a dimer in solution (Figure 4G, H), suggesting that the very C-terminus of Panx harbors an additional dimerization activity, a model consistent with observations that several LC8 client proteins harbor alternative dimerization entities directly adjacent to their LC8 binding sites^46,61^. Immediately downstream of the two TQT motifs in Panx are two predicted alpha-helices, which could mediate such additional dimerization. Truncation of these two helices alone did not prevent dimer formation of the quaternary SFiNX complex containing LC8 (Figure 4G,H), but in combination with mutation of the two TQT motifs (Panx[LC8-mut_ΔC-helix]), resulted in a monomeric, ternary complex (Figure 4G,H). Taken together, SFiNX is a homo-dimeric complex consisting of two Panx subunits, two Nxf2-Nxt1 heterodimers and two LC8 homodimers. The two C-terminal alpha-helices in Panx probably serve as an initial dimerization unit, ensuring that LC8 binding enforces homo-dimerization rather than bridging SFiNX to other LC8 client proteins.

### SFiNX dimerization is required for co-transcriptional silencing

To determine whether SFiNX dimerization is required for piRNA-guided co-transcriptional silencing we probed the ability of Panx variants, lacking either one of its two dimerization activities alone or both together, to restore transposon silencing in OSCs depleted for endogenous Panx. While deletion of the two C-terminal alpha-helices (Panx[ΔC-helix]) impacted SFiNX silencing activity only mildly, mutation of the two LC8 binding motifs (Panx[LC8-mut]) reduced it markedly. Deletion of both dimerization interfaces (Panx[ΔC-term]) resulted in complete loss of SFiNX activity (Figure 5A), indicating that dimerization is required for SFiNX function.

**Fig. 5.**
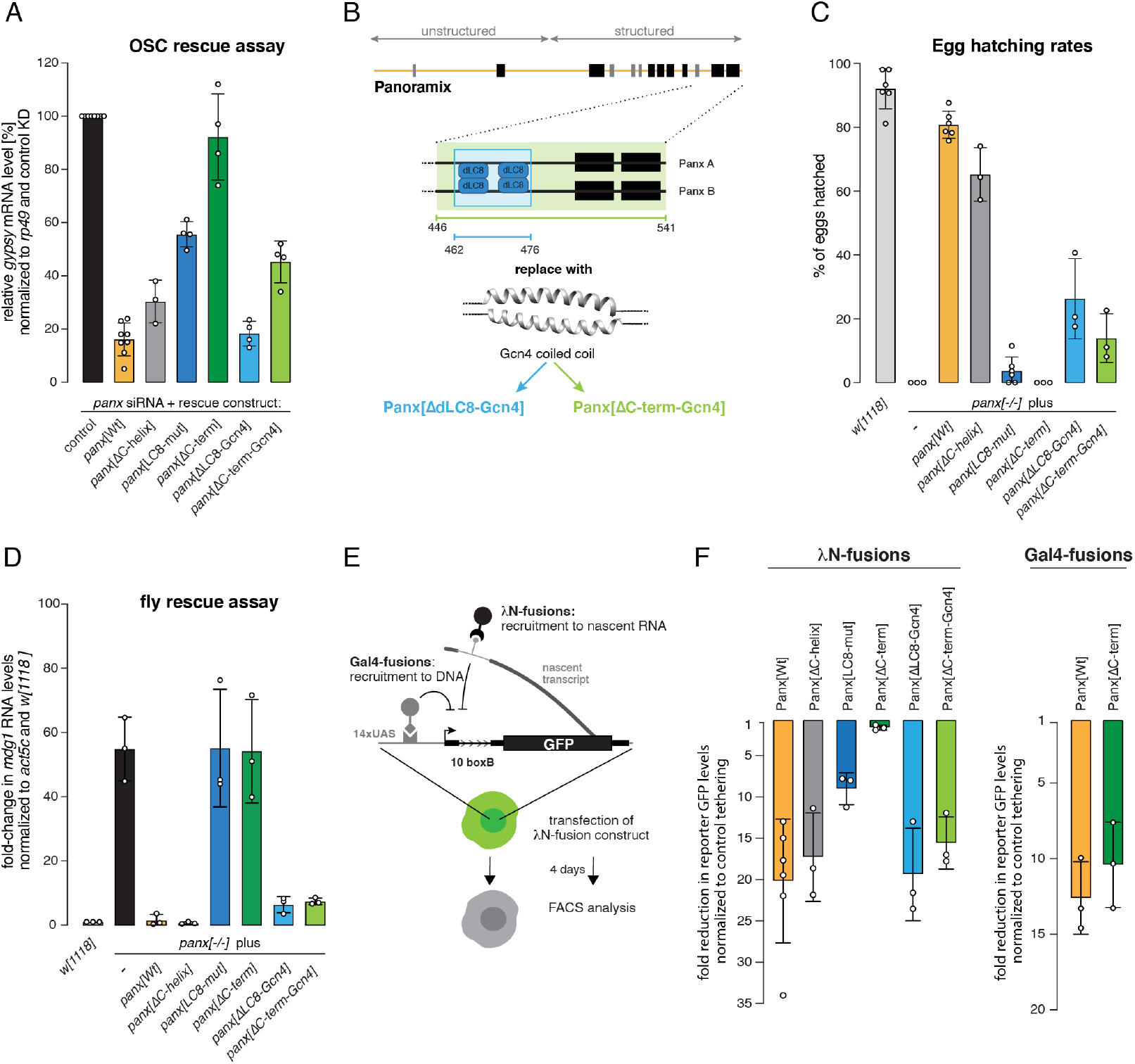
Dimerization is required for SFiNX to silence in a cotranscriptional manner. **A**, Bar graph showing the transposon repression rescue potential of indicated Panx expression constructs transfected into OSCs depleted of endogenous Panx. *gypsy* levels were determined via qRT-PCR (at least 3 biological replicates are shown; error bars: St.dev.). **B**, Cartoon representation of the ‘bypass’ strategies using the coiled-coil domain from yeast GCN4. Internal replacement of both LC8 binding sites (blue) or replacement of the entire C-terminal domain of Panx (green) are shown. **C**, Bar graph showing hatching rate of eggs laid by flies with indicated genotype (at least 3 biological replicates are shown; error bars: St.dev.). **D**, Bar graph showing mRNA levels of the LTR retrotransposon *mdg1* determined via qRT-PCR in ovaries of indicated genotypes. Expression levels were normalized to *w[1118]* controls and *act5C* (n = 3 biological replicates; error bars: St.dev.). **E**, Cartoon showing the OSC reporter system that allows analysis of co-transcriptional (λN-recruitment) and transcriptional (Gal4 recruitment) silencing. **F**, Bar diagrams showing silencing potential (GFP repression) of indicated λN-(left) or Gal4-(right) fusion proteins normalized to control (data represent mean + St.dev. of at least 3 independent experiments).

We hypothesized that, if homo-dimerization is the sole function of the Panx C-terminus, replacing it with a heterologous dimerization module should rescue SFiNX functionality. To test this, we replaced either the internal 15 amino-acid peptide harboring both LC8-binding motifs (Panx[ΔLC8-Gcn4]) or the entire Panx C-terminus (lacking both LC8 binding sites and the C-terminal helices; Panx[ΔC-term-Gcn4]) with a 32 amino-acid motif from the yeast transcription factor Gcn4 that forms a coiled coil which is sufficient to promote dimerization^62,63^ (Figure 5B). Gcn4 coiled coil mediated dimerization was confirmed using transiently expressed FLAG-tagged and GFP-tagged Panx (Figure S5A). When tested in OSC rescue experiments, both SFiNX variants harboring the Gcn4 coiled coil exhibited substantially improved transposon silencing activity compared to the dimerization-defective counterparts (Figure 5A, S5B, C).

We probed the functionality of the same Panx variants in flies. *panx* mutant females expressing tagged wildtype Panx or any of the dimerization-defective variants from a genomic transgene were analyzed for sterility and transposon silencing (Figure S5D). The results confirmed that SFiNX[Panx LC8-mut] is a strong mutant (egg hatching: ∼4%) and that monomeric SFiNX[Panx ΔC-term] is equivalent to a *panx* null mutant (the few laid eggs displayed a collapsed morphology and egg hatching was 0%) (Figure 5C). Expression of Panx variants harboring the Gcn4 coiled coil substantially restored transposon silencing and fertility compared to their respective dimerization-defective counterparts (Figure 5 D). Thus, a heterologous protein dimerization domain can largely replace the function encoded in the 95-amino acid Panx C-terminus. To determine whether dimerization of SFiNX is required for interaction with upstream or downstream factors, we used the silencing reporter assay that compares DNA-targeted versus RNA-targeted silencing (Figure 5E)^35^. We recruited the different SFiNX variants to nascent reporter RNA and tested their silencing capacity. In this assay, events upstream of SFiNX (e.g. its recruitment to the target by Piwi) are bypassed, and only the ability of SFiNX to mediate co-transcriptional silencing is assessed. Deletion of the C-terminal helices did not impair SFiNX functionality, however mutation of both LC8 binding sites led to substantially reduced activity (Figure 5F; S5E). The monomeric SFiNX (Panx[ΔC-term]) complex lost all co-transcriptional silencing capacity. Gcn4 coiled-coil mediated dimerization fully restored SFiNX silencing ability, even for the variant lacking 95 C-terminal amino acids in Panx (Figure 5F; S5E). Thus, dimerization of SFiNX is required downstream of its recruitment to a target RNA via Piwi.

To assay whether SFiNX dimerization is required for recruiting heterochromatin effectors we targeted monomeric (Panx[ΔC-term]) or dimeric, wildtype SFiNX directly to the reporter DNA using UAS-sites upstream of the promoter and fusion proteins between Panx and the Gal4 DNA binding domain. Monomeric SFiNX was as potent at transcriptional silencing as wildtype SFiNX^35^ (Figure 5F; S5E). Thus, SFiNX dimerization is not required for it to signal to the downstream silencing machinery. Taken together, we conclude that dimerization of SFiNX is required for the N-terminal silencing domain within Panx^35-38^ to silence the target locus in a cotranscriptional manner, meaning when recruited to chromatin via a nascent target RNA.

### Dimeric SFiNX interacts with nucleic acids

To understand how dimerization enables SFiNX’ co-transcriptional silencing activity, we characterized monomeric versus dimeric SFiNX complexes *in vitro*. We first assessed whether SFiNX interacts with nucleic acids using electrophoretic mobility shift assays (EMSA). We purified recombinant SFiNX lacking the N-terminal low complexity domain of Panx (involved in silencing) and the tandem RRM-LRR domains of Nxf2 (implicated in ssRNA binding) (Figure 4G)^35-38^. At physiological salt and Mg^2+^ levels, dimeric SFiNX interacted with nucleic acids, preferentially binding single-stranded (ss) RNA, having lower yet comparable affinities for double-stranded (ds) DNA and dsRNA, and binding ssDNA only weakly (Figure 6A,B). Deletion of the Panx C-terminal helices or mutating both LC8 binding sites substantially weakened nucleic acid binding (as measured for ssRNA) (Figure 6C, S6A). Simultaneous deletion of both dimerization interfaces abolished RNA binding (Figure 6C, S6A). Thus, SFiNX binds to nucleic acids in a dimer-dependent manner.

**Fig. 6.**
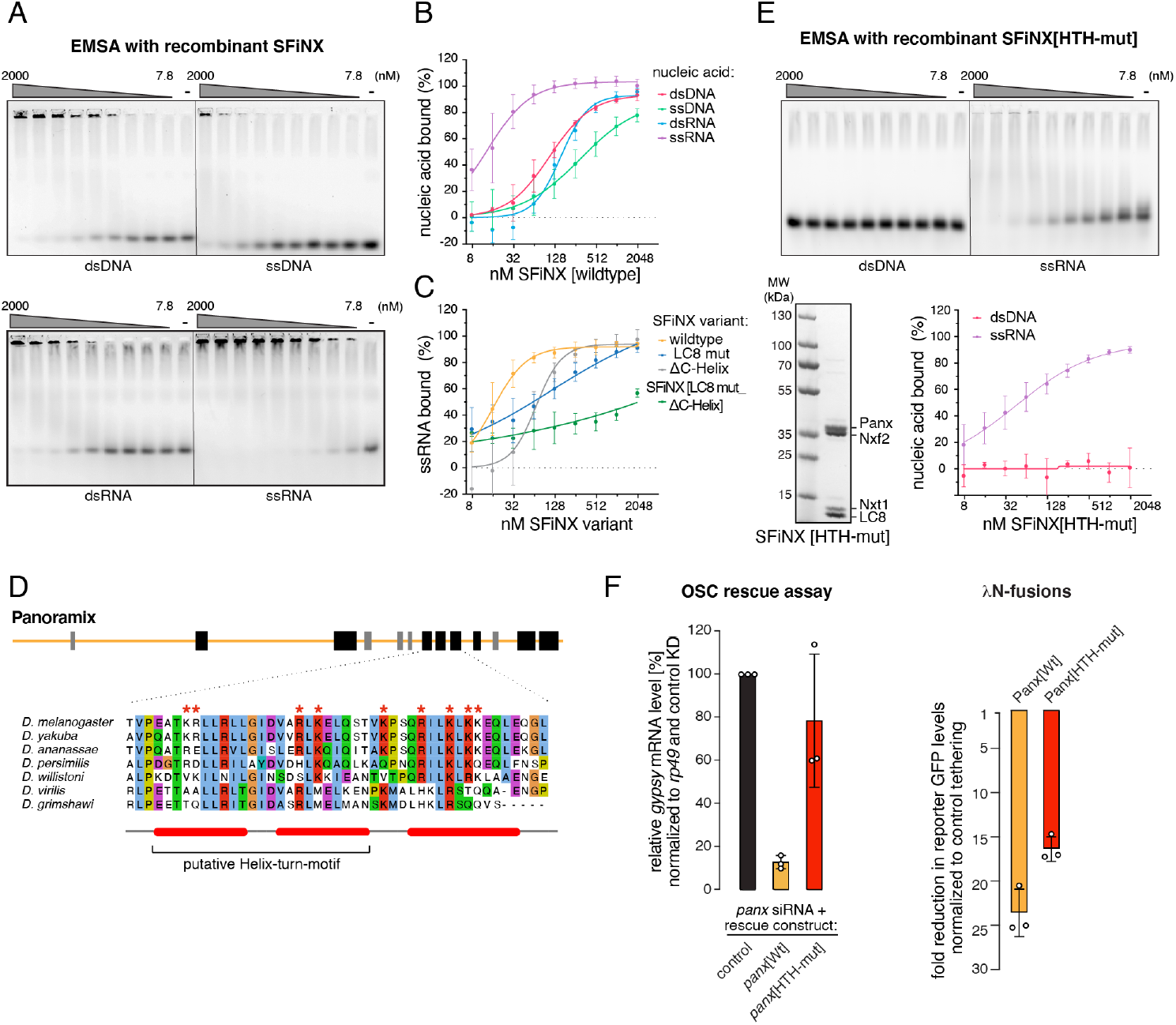
SFiNX interacts with nucleic acids. **A**, Agarose gel images showing electrophoresis mobility shift assays (EMSA) using different types of nucleic acids (Alexa-647 labeled). Serial dilution of recombinant SFiNX[wildtype] ranging from 7.8 to 2000 nM is shown (‘-’ indicates no protein; nucleic acid concentration: 5nM). **B**, Binding curves based on EMSA experiments in A (calculated from quantified free nucleic acid; n=3; error bars: St.dev.) are shown. **C**, Binding curves between ssRNA and indicated SFiNX variants are shown (n=3; error bars: St.dev.). **D**, Schematic representation of Panx, indicating predicted secondary structure elements (black: alpha-helices, grey: beta-strands) and the putative HTH-motive with its evolutionary conservation below. Residues mutated in E and F are labeled with asterisks. **E**, To the top, agarose gel images showing electrophoresis mobility shift assays (EMSA) using different types of nucleic acids (Alexa-647 labeled). Serial dilution of recombinant SFiNX[wildtype] ranging from 7.8 to 2000 nM is shown (‘-’ indicates no protein; nucleic acid concentration: 5nM). Coomassie stained SDS PAGE showing recombinant SFiNX complex with HTH mutations (bottom left). Binding curves based on EMSA experiments with SFiNX[HTH-mut] with ssRNA and dsDNA (calculated from quantified free nucleic acid; n=3; error bars: St.dev.) are shown. **F**, To the left, bar graph showing the transposon repression rescue potential of indicated Panx expression constructs transfected into OSCs depleted of endogenous Panx. *gypsy* levels were determined via qRT-PCR (n=3 biological replicates are shown; error bars: St.dev.). To the right, bar diagrams showing silencing potential (GFP repression) of indicated λN-fusion proteins normalized to control (data represent mean + St.dev. of 3 independent experiments).

The interaction of SFiNX with dsDNA was intriguing as this suggested a putative model where SFiNX enables or stimulates cotranscriptional silencing by tethering the nascent target RNA (via an interaction with Piwi) to the underlying chromatin locus. To test this, we set out to identify mutations in SFiNX that are compatible with dimer formation yet defective in DNA binding. Upstream of the LC8 binding sites, Panx harbors an alpha-helical region that is rich in positively charged residues and that exhibits remote similarity to a helix-turn-helix motif, a widespread domain that binds DNA. Mutation of nine positively charged amino acids in this region (Figure 6D) of Panx resulted in a recombinant SFiNX complex that, in respect to dimer formation, behaved as the wildtype complex, but that was incapable of binding to dsDNA and showed weaker binding to ssRNA (Figure 6E, S6B). Genetic rescue experiments indicated that this DNA-binding defective Panx mutant is strongly impaired in compensating for endogenous Panx in OSCs (Figure 6F, S6C). When recruited to the nascent reporter RNA via the λN-boxB system, DNA binding deficient Panx was instead only moderately impaired in its ability to support co-transcriptional silencing (Figure 6F, S6D). Our findings indicate that SFiNX’ ability to bind DNA (or ssRNA) is important for its *in vivo* function, but that this is not the central feature that enables SFiNX to elicit cotranscriptional silencing.

### SFiNX forms molecular condensates in a nuclear acid stimulated manner

Many proteins functionally implicated in chromatin processes have the ability to form molecular condensates, often in a nucleic acid stimulated manner. Intrigued by the observation that recombinant SFiNX-nucleic acid complexes hardly entered the gel in our EMSA experiments, we set out to investigate whether SFiNX is able to form molecular condensates *in vitro*.

When lowering the salt concentration of a 10 micromolar SFiNX solution to physiological levels (150 mM NaCl), the previously clear solution rapidly became turbid due to the formation of phase-separated droplets (Figure 7A). To characterize this process, we purified a GFP-tagged SFiNX variant and mixed it with untagged recombinant SFiNX at a ratio of one to ten. Imaging the phase separation behavior using microscopy revealed that SFiNX forms molecular condensates in a concentration dependent manner with a threshold concentration of ∼7 micromolar (Figure 7A, B). The condensates were round and occasionally fused into larger round condensates, suggesting that they have liquid characteristics^64^ (Figure 7C). We analyzed the tagged and untagged SFiNX mixture using fluorescence recovery after photobleaching (FRAP) and observed a fast and complete recovery of fluorescence (Figure 7D) confirming that SFiNX forms liquid condensates *in vitro* under physiological salt concentrations.

**Fig. 7.**
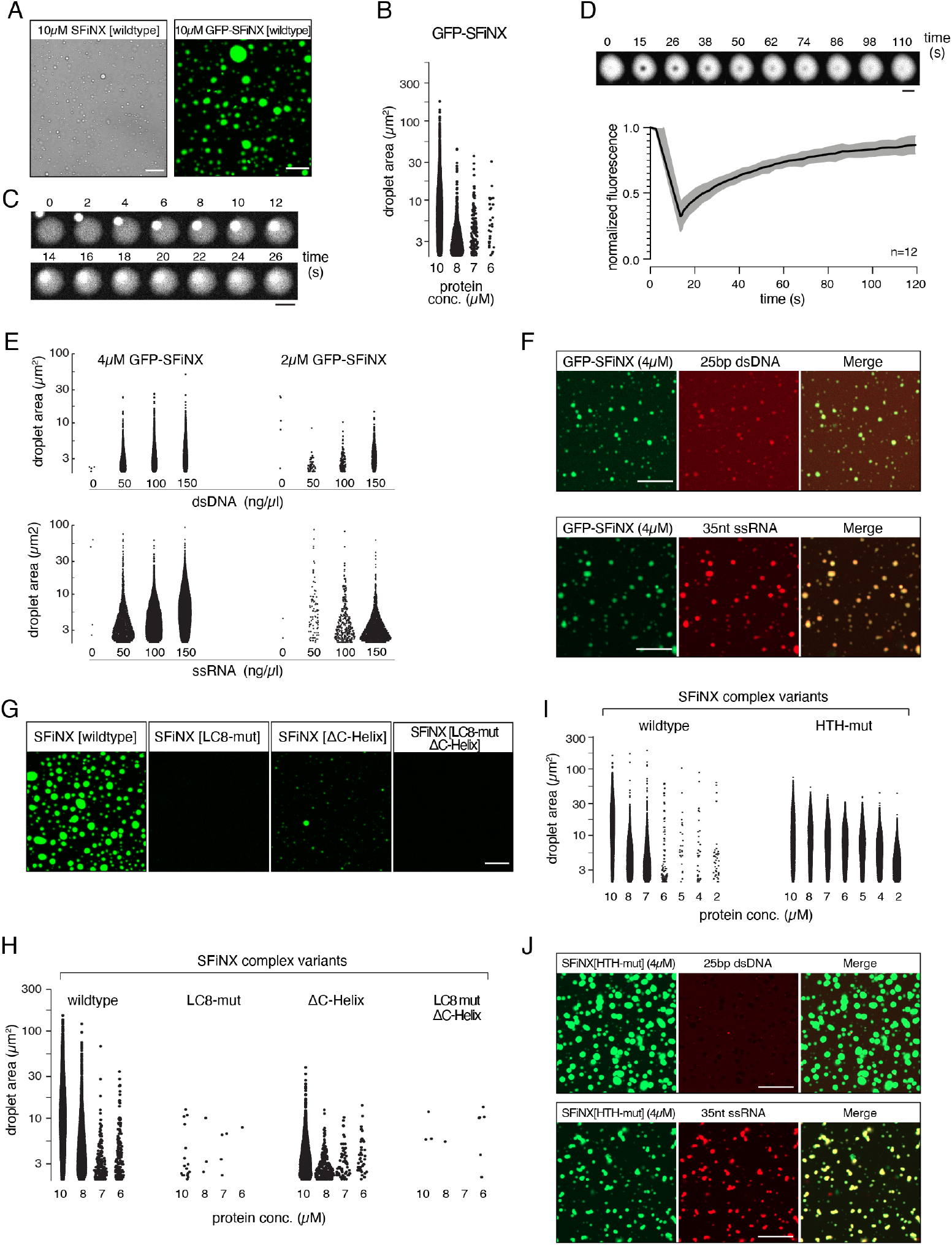
SFiNX forms molecular condensates, stimulated by nucleic acid binding. **A**, Bright-field image showing SFiNX[Wt] forms condensates *in vitro* in the absence at 10μM (left, scale bar: 20µm). Confocal image showing *in vitro* condensate formation of recombinant SFiNX complex at 10µM concentration (containing 10% mEGFP tagged SFiNX; scale bar: 20µm). **B**, Shown are droplet number and area per droplet (sum of three microscopic fields of view) of recombinant SFiNX condensates (with mEGFP-SFiNX at 1:10) at indicated protein concentration. **C**, Time lapse series showing fusion event of two SFiNX condensates (differing intensities are due to the bigger droplet being photo-bleached prior to the fusion event; scale bar: 5µm). **D**, Time lapse series showing recovery after partial photo bleaching of SFiNX condensates (confocal image: representative experiment, scale bar: 5µm; below: quantification of FRAP data; n=12 experiments; St.dev. in gray). **E**, Shown are droplet number and area per droplet (sum of three microscopic fields of view) of recombinant SFiNX condensates (with mEGFP-SFiNX at 1:10) at indicated protein and dsDNA (top) ssRNA (bottom) concentrations. **F**, Confocal images (scale bar: 20µm) showing enrichment of Alexa647 labeled double stranded DNA (top) or single stranded RNA (bottom) in condensates of mEGFP-SFiNX (4µM). **G**, Confocal images showing ability of condensate formation of indicated, fluorescein-labeled SFiNX variants (concentration: 10µM; scale bar: 20µm). **H**, Shown are droplet number and area per droplet of three microscopic fields of fluorescein-labeled SFiNX variants at indicated protein concentration (representative images: Figure 7G, S7C). **I**, Shown are droplet number and area per droplet of three microscopic fields of fluorescein-labeled SFiNX variants at indicated protein concentration (representative images: Figure S7D). **J**, Confocal images (scale bar: 20µm) showing enrichment (ssRNA) or exclusion (dsDNA) of Alexa647 labeled nucleic acids in condensates of fluorescein-labeled SFiNX[HTH-mut] (4µM).

Considering SFiNX’ ability to bind to nucleic acids (Figure 6), we investigated whether presence of ssRNA or dsDNA impacts its liquid-liquid phase separation behavior. Both, ssRNA as well as dsDNA substantially stimulated the formation of SFiNX condensates (Figure 7E, S7A, S7B), lowering the saturation concentration of SFiNX to 2 micromolar (versus ∼7 micromolar for the wildtype complex). Consistent with this and with the previous EMSA experiments, labelled ssRNA or dsDNA co-partitioned into SFiNX condensates (Figure 7F). In conclusion, the recombinant SFiNX complex used in our experiments, though being nearly entirely composed of structured domains, exhibits a pronounced ability to form molecular condensates in a nucleic acid stimulated manner.

If formation of molecular condensates underlies SFiNX’ ability to enable co-transcriptional silencing, monomeric SFiNX, which is incapable of co-transcriptional silencing (Figure 5), should be defective in condensate formation. We therefore studied the phase separation properties of recombinant SFiNX variants lacking either the C-terminal helices, the dimerization protein LC8, or both (Figure 7G, H, S7C). In order to visualize and quantify droplet formation, we added 5% of fluorescein-labeled complexes to unlabeled complexes. Phase separation behavior of wildtype SFiNX was comparable to the one of the GFP-tagged complex. In contrast, SFiNX complexes with weakened dimerization ability showed a moderate (Panx[ΔC-helix]) to severe (Panx[LC8-mut]) reduction in condensate formation. Monomeric SFiNX (Panx[LC8-mut_ΔC-helix]) was unable to form condensates. Thus, the ability of SFiNX to form molecular condensates, just as its co-transcriptional silencing capacity, requires its dimerization. We note that the strength of the *in vivo* phenotype observed for the different SFiNX dimerization mutants correlated precisely with their phase separation ability *in vitro* (wildtype > Panx[ΔC-helix] > Panx[LC8-mut] > Panx[LC8-mut_ΔC-helix]). This argues that the ability to form molecular condensates is central for SFiNX’ co-transcriptional silencing activity. It also predicts that the dimeric yet DNA-binding defective SFiNX complex, which is capable of co-transcriptional silencing in the tethering assay (Figure 6E, F) should still be capable of condensate formation. Indeed, the SFiNX variant containing Panx[HTH-mutant] exhibited robust phase separation behavior *in vitro* with a saturation concentration five times lower than that of the wildtype complex (Figure 7I, S7D). Consistent with the EMSA results, dsDNA was excluded from the condensates (Figure 7J), supporting the inability of this SFiNX variant to interact with DNA. Taken altogether, our data strongly point to the critical importance of multi-valent interactions within the SFiNX complex that *in vitro* underlie its phase separation behavior and that *in vivo* are required for empowering the unstructured, N-terminal Panx domain to establish heterochromatin in a co-transcriptional manner.

## Discussion

In this work, we show that SFiNX, a key protein complex involved in Piwi-mediated heterochromatin formation, is able to form molecular condensates in a dimer-dependent manner. The ability to form condensates correlates precisely with SFiNX’ ability to establish a robust co-transcriptional silencing response when the complex is targeted to nascent RNA. Previous work has shown that the N-terminal unstructured portion of the Panx subunit within SFiNX, while necessary for silencing, is not sufficient to induce silencing when recruited to nascent RNA^35,37^. Our experiments demonstrate that SFiNX forms multivalent interactions in a nucleic acid stimulated manner suggesting this process is involved in the formation of molecular condensates at actively transcribed chromatin. The saturation concentration for condensate formation in our experiments was ∼7 µM and it is lowered to ∼2 µM in the presence of nucleic acids. These critical concentrations are considerably lower than those where HP1 forms condensates (∼40 µM)^65^ and are similar to those where transcriptional regulators like the pioneer factor Klf4 show phase separation behavior (Morin et al. 2020; biorXiv). Our mutational studies show that the ability of SFiNX to form condensates *in vitro* correlates precisely with its ability to induce heterochromatin formation in a co-transcriptional manner *in vivo*. We demonstrate that SFiNX multivalency is required downstream of its recruitment to the target transcript through Piwi, and that it is dispensable for SFiNX to signal to the heterochromatin machinery. This argues that SFiNX condensate formation, stimulated by nucleic acid binding, enables co-recruited silencing activities to efficiently modify the underlying chromatin locus, potentially by increasing the dwell time of the nascent target RNA at the site of transcription.

The nuclear SFiNX concentration is expected to be considerably lower than the saturation concentration of ∼7 μM at which condensate formation is observed *in vitro*. Consistent with this, microscopically detectable SFiNX condensates are not seen in wildtype cells. However, transposon transcripts harbor hundreds of complementary piRNA targets, which would recruit multiple copies of SFiNX via Piwi, to the site of transcription, thereby elevating SFiNX concentration locally. We propose that this allows for multivalent interactions of SFiNX complexes and condensate formation at piRNA target sites in a process that would be further stimulated by nascent RNA and the underlying DNA locus. Such a model would explain why single piRNA target sites are insufficient to induce co-transcriptional silencing^66^, and why increasing the number of Piwi or SFiNX target sites in a reporter gene leads to highly synergistic silencing responses^30,66^.

Also in fission yeast, oligomerization of the RNA induced transcriptional silencing (RITS) complex is critically involved in small RNA-guided heterochromatin formation^67^. RITS consists of siRNA-loaded Ago1, Tas3, and Chp1^67^ and binds target RNA via Ago1 to recruit chromatin-modifying factors^13-15^. Two additional activities are required for silencing. First, Chp1 harbors a chromodomain that binds the H3K9 methyl mark^68^. The RITS complex can therefore tether the nascent target RNA to the underlying chromatin locus once it has been modified by the H3K9 methyl-transferase complex. Second, Tas3 mediates RITS complex oligomerization via its C-terminal, alpha-helical domain^69^, which is required for the silencing response to spread at the target locus. These s and tethering of the target RNA at chromatin is a general phenomenon in small RNA-guided co-transcriptional silencing processes. Liquid-liquid phase separation and RNA retention at chromatin has also been implicated in mammalian X-chromosome inactivation^70,71^ and in the function of long non-coding RNAs at chromatin^72^. Our work now links small RNA guided cotranscriptional gene silencing to the formation of condensates adding to the emerging theme that molecular condensates are critically involved in gene regulatory and chromatin-related processes.

We have identified the dynein light chain LC8 as a subunit of the SFiNX silencing complex. LC8 is a highly conserved protein in animals (96.6% identity between flies and humans) and forms a homodimer, enabling the dimerization of a large variety of proteins and protein complexes^46,47^. Our data further support the view that LC8 mediates homo-dimerization of protein complexes rather than bridging different client proteins in hetero-dimeric complexes. Intrinsic dimerization activities flanking LC8 binding sites (like the C-terminal alpha-helices in Panx) are probably involved in promoting homo-dimeric assemblies, which are then further stabilized through LC8 binding. This would prevent the erroneous formation of non-functional heterodimers. In the case of SFiNX, homodimerization strictly correlates with its ability to form molecular condensates and to bind nuclei acids. Besides SFiNX, we show that several other nuclear protein complexes are LC8 clients and hence act as homodimers. While the precise molecular role of dimerization in other LC8 client complexes may differ, it is possible that stimulating nucleic acid binding and enabling condensate formation by promoting multivalent interactions of protein complexes may be common to many nuclear molecular machines.

## Author Contributions

J.S. and J.Batki made the initial discovery of LC8 as a SFiNX member. J.S. performed all molecular biology and fly experiments, except: U.H. performed the SEC-MALS experiments of insect cell complexes, M.G. performed Cut & Run experiments and analysis, A.I.V. performed co-IP of Panx, K.P. mapped the LC8 interaction sites in Woc and Tlk. J.W. generated, purified and grew crystals of the LC8-Panx complex from bacteria, performed the X-ray crystallographic analyses, and performed the SEC-MALS assay and the ITC measurements under the supervision of D.J.P. Tagged mES cells were generated by N.F., all fly stocks were generated by P.D. M.N. generated the phylogenetic comparisons. K.M. supervised mass spectrometry analysis. The project was supervised by J.B., D.J.P. and C.P. The paper was written by J.S. and J.B. with input from all authors. The authors declare no conflict of interest.

## Acknowledgments

We thank K. Meixner for experimental support, and A. Meinhart and A. Vogel for advice on the biochemical characterization of SFiNX. We thank S. Schweighofer for advice on image analysis and the Siomi lab for the Eggless antibody. The following VBCF core facilities provided outstanding support: Protein Technologies Facility (protein expression), NGS facility (Illumina sequencing), VDRC (fly stocks). We thank the IMBA Stem Cell Core Facility for generating tagged mouse ES cells and the GMI/IMBA/IMP Bio-optics unit for support with confocal imaging. We thank G. Riddihough from Life Science Editors for comments on the manuscript (http://lifescienceeditors.com), the Brennecke lab, in particular L. Baumgartner, D. Handler, C. Yu and K. Senti for support and feedback, as well as D. Gerlich and L. Cochella for feedback on the manuscript and S. Saha for discussions. The Brennecke lab is supported by the Austrian Academy of Sciences, the European Community (ERC-2015-CoG − 682181), and the Austrian Science Fund (F 4303 and W1207). X-ray diffraction studies were conducted at the Advanced Photon Source on the Northeastern Collaborative Access Team beamlines, supported by NIGMS grant P30 GM124165 and U.S. Department of Energy grant DE-AC02-06CH11357. The Eiger 16M detector on 24-ID-E beamline is funded by a NIH-ORIP HEI grant (S10OD021527). The ITC experiments were performed in the Serganov laboratory (NYU) with the help of A. Kaushik. This work was supported in part by the Maloris Foundation (DJP). The Memorial Sloan Kettering Cancer Center structural biology core facility is supported by National Cancer Institute Core grant P30-CA008748. This work of K.M. was financed by the EPIC-XS, Project Number 823839 and the Horizon 2020 Program of the European Union and the by the ERA-CAPS I 3686 project of the Austrian Science Fund. U.H. was supported by the SNF Early Postdoc Mobility fellowship (P2GEP3_188343). M. Gehre was supported by the VIP2 Post-Doc fellowship program as part of the EU Horizon 2020 research and innovation program (Marie Skłodowska-Curie grant No. 847548).

## Materials and Methods

### Fly strains

All fly strains used in this study are listed in Table S3 and are available from the VDRC <u>(h</u>t<u>tp://stockcenter.vdrc.at/control/main).</u> Flies were kept at 25 °C and for each experiment, flies were aged for 4 days and kept on apple juice agar plates with yeast paste to ensure consistent ovarian morphology. Flies with an extra genomic copy of GFP-tagged LC8^74^ were generated by recombination-mediated genetic engineering as described in^75^. Rescue strains with different Panx variants were generated as described in^35^. Point mutations in the LC8 interaction sites of the endogenous Panx locus were generated as described in^76^ using an HDR donor oligo and guide RNAs listed in Table S4. Guide RNAs were cloned into the pCFD4 plasmid via PCR followed by Gibson assembly.

### OSC cell culture

OSCs were cultured as described^39,40^. Plasmid and siRNA transfections were performed using Cell Line Nucleofector kit V (Amaxa Biosystems) with the program T-029, using 6 million cells per transfection. siRNAs used in this study are listed in Table S5.

### Generation of endogenously tagged OSC cell lines

The homology directed repair (HDR) template for N-terminal tagging of LC8 and Eggless consisted of a Puromycin-resistance followed by a P2A cleavage site followed by a FLAG-GFP-tag. The tagging construct was flanked by approximately 500bp homology arms around the start codon of the respective gene and was cloned into the pBluescriptII SK (+) vector. This plasmid was used as a template for PCR and 2500ng of purified HDR template PCR product combined with 1500ng of guide RNA expression plasmid (Addgene 49330) containing the guide RNA listed in Table S4 were co-transfected into OSCs. After two days, cells were plated in different dilutions and on the following day Puromycin containing medium (5µg/ml) was added to select for individual clones for 4 days. After additional 7 days of growth individual clones were picked and expanded. Successful tagging was analyzed by PCR, FACS and western blot. For C-terminal tagging of Panx, a GFP-FLAG tag followed by a Puromycin-resistance gene driven by the *traffic jam* enhancer in combination with the *Drosophila* synthetic core promoter (DSCP)^77^ was used. The HDR tagging was performed as described above.

### Co-immunoprecipitation of endogenously tagged proteins

OSC/mES cells were collected after trypsinization by centrifugation and washed with PBS. For isolation of nuclei the cell pellet was resuspended in Buffer 1 (10 mM Tris-HCl pH=7.5, 2 mM MgCl2, 3 mM CaCl2, supplemented with Roche Complete Protease Inhibitor Cocktail), incubated at 4°C for 20 min followed by a centrifugation step. The pellet was resuspended in Buffer 2 (10 mM Tris-HCl pH=7.5, 2 mM MgCl2, 3 mM CaCl2, 0.5 % IGEPAL CA-630, 10 % glycerol, supplemented with Roche Complete Protease Inhibitor Cocktail), incubated at 4°C for 10 min followed by a centrifugation step. The isolated nuclei were lysed in Buffer 3 (20 mM HEPES pH=7.5, 150 mM NaCl, 2 mM MgCl2, 0.3 % Triton X-100, 0.25 % IGEPAL CA-630, 10 % glycerol, supplemented with Roche Complete Protease Inhibitor Cocktail), incubated at 4°C for 20 min followed by sonication using Diagenode Bioruptor for 10 min (30 sec ON, 30 sec OFF) at low intensity. Lysate was cleared by centrifugation and incubated for 2h at 4°C with magnetic GFP-Trap agarose (Chromotek). The beads were washed 3 x 10 min with Buffer 3 and were either used for mass spectrometry analysis or the proteins were eluted in 1× SDS buffer with 5 min incubation at 95°C for western blotting. For co-immunoprecipitation of LC8 from OSCs and ovaries, isolation of nuclei was omitted, and cells were directly lysed in Buffer 3 and processed as described above.

### Silver staining of SDS PAGE gels

4-20% precast SDS PAGE gels (BioRad) were used for electrophoretic separation of immuno purified samples. Silver staining was carried out using the Pierce Silver Stain Kit (Thermo Fisher Scientific) according to manufacturer’s instructions.

### Western blot

Western blot was carried out as described in^35^. For LC8/Dynll1 immunoblotting, samples were separated using 15% SDS PAGE at constant voltage. After equilibration of gels in SDS-free transfer buffer (50 mM Tris base (Trizma), 384 mM glycine) for 10 minutes, proteins were transferred onto 0.2µm PVDF membrane at 25V for 16h. After blocking the membrane with 4% milk in PBS containing 0.2% Tween-20 for 30min, antibody diluted in blocking solution was added for 4h at room temperature. The membranes were washed 3 x for 10 min with 1x PBST (0.2% Tween-20 in PBS) followed by incubation with HRP coupled secondary antibody diluted in blocking buffer for 1h. After incubation with Clarity Western ECL Substrate (BioRad) blots were imaged using a ChemiDoc acquisition system (BioRad). Antibodies are listed in Table S6.

### Immunofluorescence staining of ovaries

Immunofluorescence staining was performed as described in^35^. The mounted samples were imaged with a Zeiss LSM-780 confocal microscope. Antibodies are listed in Table S6.

### Tethering reporter assay

Tethering reporter assays were performed as previously described^35^. Fold repression compared to control experiments was calculated using 2500 cells per experiment (GFP intensity was measured with FACS). For tethering of Ctp and Nxf2 in combination with siRNA knockdown, siRNAs were first transfected and after 2 days a second transfection with siRNA and tethering plasmid was carried out. Western blots to confirm depletion of proteins and FACS to monitor GFP expression were performed on day 5.

### RT-qPCR

RT-qPCR was performed as previously described^35^. qPCR primers are listed in Table S4.

### Fertility measurements

For fertility assays, virgins of the respective genotype were collected and mated with w[1118] males. After aging for 4 days, 5-10 females were put in cages with apple juice agar plates containing yeast paste and allowed to lay eggs for 15h. Hatching rates of the eggs was determined 24h after removal of flies by manual scoring.

### Cut and Run

Cut&Run was performed according to instructions in CUT&RUN protocol V.3^78^ with minor modifications. In brief, 500,000 OSCs were bound to 10 µl Concanavalin A-coated magnetic beads (Polysciences, #86057-3) and lysed using Dig-wash buffer (20 mM HEPES pH 7.3, 150 mM NaCl, 0.5 mM spermidine, 0.01% digitonin, Roche Complete Protease Inhibitor -EDTA). Bead-bound cells were incubated with 0.5 µg of respective antibody (listed in Table S6) at 4°C overnight on a nutator. Afterwards, cells were washed twice, resuspended in Dig-wash buffer containing 700 ng/ml pAG-MNase (produced in house) and incubated for 1 hour at 4°C on a nutator. Cells were washed twice and resuspended in Dig-wash buffer containing 2 mM Ca^2+^ to activate pAG-MNase. Reaction was stopped by the addition of 2x STOP Buffer, and samples were incubated at 37°C mixing at 500 rpm for 15 minutes to release DNA fragments into solution. After centrifugation, 0.1% SDS and 0.2 μg/μl Proteinase K were added to the supernatant and samples were incubated for 1 hour at 55°C. DNA was purified using a DNA purification kit and libraries were prepared following the manufacturer’s instructions with NEBNext Ultra II DNA Library Prep Kit for Illumina. Sequencing was performed on a HiSeqV4 using 50 bp single-end mode.

### Data Analysis

For whole genome analysis of Cut&Run data, sequencing reads were aligned to the fly reference genome (dm6 assembly) using Bowtie2 (Galaxy v. 2.3.4.3)^79^ with zero mismatches allowed. Only nonduplicated, uniquely mapped reads were retained for further analysis. The plots to visualize the distribution of H3K9me3 around euchromatic transposon insertion sites^31^ was generated using ngs.plot (v.2.61)^80^.

### mES cell culture

An3-12 mES cells were used for all experiments and were cultivated as described^81^.

### Endogenous tagging of mouse ES cells

The Homology directed repair (HDR) template for N-terminal tagging of Dynll1 contained the epitope recognized by the BC2-nanobody followed by a triple FLAG tag and an AID tag. This tag was flanked by approximately 500bp long homology arms flanking the start codon of the Dynll1 gene. The HDR template was cloned into the TOPO backbone and guide RNAs were cloned into the guide RNA expression plasmid (Addgene 71707) containing an mCherry expression cassette. Plasmids were co-transfected into An3-12 mES cells stably expressing Tir1 from the Rosa26 locus using Lipofectamine 2000. After 2 days, mCherry expressing cells were FACS sorted and sparsely plated to allow growth of individual clones. After additional 7 days of growth, individual clones were picked and characterized with PCR and western blot. Guide RNA sequences are listed in Table S4.

### Mass spectrometry

Mass spectrometry was carried out as described in^35^.

### Enrichment analysis of LC8 binding sites in co-immunoprecipitation experiments

Enrichment analysis for LC8 binding motives was done by Gene set enrichment analysis (GSEA)^82^ using the fold enrichment of the identified proteins in the indicated co-immunoprecipitation experiments as the rank metric score and LC8 hub^45^ to predict the LC8 binding motives.

### Protein sequence alignments

Protein sequence alignments were carried out as described in^35^. For mapping orthologous genes of *Drosophila melanogaster* to *Mus musculus*, ortholog assignment of Flybase was used (FB2018_06). Only orthologs which were supported by more than 5 methods were included allowing one mouse entry to map to multiple fly entries.

### Immunofluorescence of mES cells

mES cells were seeded on Concanavalin A treated coverslips and fixed after 6h for 10min with 4% PFA in PBS at RT. After 3x washing with PBS the cells were permeabilized with 0.5% TritonX-100 in PBS for 10min followed by blocking with blocking buffer (1% BSA, 0.01% Triton X-100 in PBS) for 30 min. Cells were incubated with primary antibody diluted in blocking buffer over night at 4°C. After washing with PBS, fluorophore conjugated secondary antibody diluted in blocking buffer was added for 2 hours. Cells were washed with PBS 4 x and DNA was stained using DAPI. Confocal images were acquired on a Zeiss LSM-780 confocal microscope and the images were processed using FIJI/ImageJ. Antibodies are listed in Table S6.

### Protein expression and purification

For crystallization, *Drosophila melanogaster* LC8 (residues 1-89, LC8), as well as Panx peptides (residues 455-466 (Panx1), 467-480 (Panx2) and 455-480 (Panx1+2)) were cloned into a modified RSFduet-1 vector (Novagen) with an N-terminal His6-SUMO tag on the Panx peptides and no tag on LC8. Panx and LC8 were co-expressed in *E. coli* strain BL21(DE3) RIL (Stratagene). The cells were grown at 37°C until OD600 reached 0.8, then the media was cooled to 16 °C and IPTG was added to a final concentration of 0.35 mM to induce protein expression overnight at 16 °C. The cells were harvested by centrifugation at 4 °C and disrupted by sonication in Binding buffer (20 mM Tris-HCl pH 8.0, 500 mM NaCl, 20 mM imidazole) supplemented with 1 mM PMSF (phenylmethylsulfonyl fluoride) and 3 mM β-mercaptoethanol. After centrifugation, the supernatant containing complexes of Panx and LC8 was loaded onto 5 ml HisTrap Fastflow column (GE Healthcare). After extensive washing with Binding buffer, the complex was eluted with Binding buffer supplemented with 500 mM imidazole. The His6-SUMO tag was removed by Ulp1 protease digestion during dialysis against Binding buffer and separated by reloading onto a HisTrap column. The flow-through fraction was further purified by HiTrap Q FF column and a Superdex 75 16/60 column (GE Healthcare). The pooled fractions were concentrated to 15 mg/ml in crystallization buffer (20 mM Tris-HCl pH 7.5, 200 mM NaCl, 1 mM DTT). The Panx mutants were generated by site-directed mutagenesis with QuickChange kit (Agilent) and confirmed by sequencing. To carry out ITC experiments, the Panx peptides and LC8 carrying a His6-SUMO tag were expressed in isolation and purified otherwise as described above.

Co-expression of protein complexes in insect cells was carried out as previously described^35^. An N-terminal His6-tag was fused to LC8 (or to Panx in case the respective complex didn’t contain LC8). Cell pellets were resuspended in Lysis buffer (50 mM Tris pH 8.0, 300 mM NaCl, 2 mM DTT, 0.3 % TritonX-100, Roche Complete protease Inhibitor Cocktail, Benzonase) followed by sonication for 10 min (1sec ON, 2ecs OFF) at 10% intensity using a Branson 450 Digital Sonifier. The lysates were cleared by centrifugation and loaded onto a 5 ml HisTrap HP column (GE Healthcare). After washing with buffer A (50 mM Tris pH 8.0, 300 mM NaCl, 2 mM DTT) the bound proteins were eluted through a gradient elution targeting 500mM Imidazole in buffer A. Peak fractions were pooled and diluted to 120 mM NaCl using buffer A with 0 mM NaCl. The protein was then loaded on a HiTrapQ HP anion exchange column (GE Healthcare) and eluted in a gradient from 120–600 mM NaCl. Fractions containing the proteins of interest were pooled and concentrated using Amicon Ultra-15 30K spin concentrators (Merck Millipore) before further purification using a HiLoad 16/600 Superdex 200 prep grade size exclusion column (GE Healthcare) in storage buffer (40 mM HEPES pH 7.9, 300 mM NaCl, 1 mM TCEP, 5 % glycerol). The pooled fractions of interest were concentrated using Amicon Ultra-15 30K spin concentrators (Merck Millipore) to ∼8 mg/ml, aliquoted, flash frozen and stored at −80°C until usage.

### Crystallization, data collection and structure determination

Crystals of Panx-LC8 complexes were grown from different solutions (Panx1+2-LC8: 0.2 M Li2SO4, 0.1 M Tris pH 8.5, 40% (v/v) PEG 400; Panx1-LC8: 0.1 M NH4Ac, 0.1 M Bis-Tris pH 5.5, 17% (w/v) PEG 10000; Panx2-LC8: 0.1 M KSCN, 30% (w/v) PEG 2000 MME) using the hanging drop vapor diffusion method at 20 °C. For data collection, the crystals were flash frozen (100 K) and collected on NE-CAT beam lines 24ID-C and 24ID-E at the Advanced Photo Source (APS), Argonne National Laboratory. The diffraction data were processed with iMosflm^83^ or the NECAT RAPD online server^84^. The structures of Panx-LC8 complexes were solved by the molecular replacement method in PHENIX^85^ using the structure of LC8 in complex with Nek9 peptide (PDB 3ZKE)^60^. The resulting model was completed manually using COOT^86^ and PHENIX refinement^85^. The statistics of the diffraction and refinement data are summarized in Table S2. All molecular graphics were generated with the PyMOL program (https://pymol.org/2/).

### Size exclusion chromatography with in-line multi-angle light scattering (SEC-MALS)

SEC-MALS experiments were performed by using an Äkta-MALS system. For the analysis of LC8 in complex with Panx peptides, proteins (500 μl) at a concentration of 1 mg/ml were loaded on a Superdex 75 10/300 GL column (GE Healthcare) and eluted with HEPES buffer (20 mM HEPES, pH 7.5, 200 mM NaCl) at a flow rate of 0.2 ml/min. SFiNX complex variants were injected at 2 mg/ml (40 μl) on a Superdex 200 increase 3.2/300 column and eluted in storage buffer without glycerol (40 mM HEPES pH 7.9, 300 mM NaCl, 1 mM TCEP) at a flow rate of 0.07 ml/min. Light scattering was monitored on a miniDAWN TREOS system (Wyatt Technologies). Molecular masses of proteins were analyzed using the Astra program (Wyatt Technologies).

### Isothermal Titration Calorimetry (ITC)

All the ITC-based binding experiments were performed on a MicroCal ITC200 calorimeter at 20°C. Panx peptides and LC8 were purified in the same buffer (20 mM Tris pH 7.5, 150 mM NaCl, 2 mM β-mercaptoethanol). Panx peptides at the concentration of 200 µM (Syringe) were titrated in to 15 µM LC8 protein in the cell. The exothermic heat of reaction was measured by sequential injections of 2 µL the peptides into protein solution with 180 s interval spacing. The data was fitted using the program Origin with one site model.

### OSC rescue assay

OSC rescue assays were carried out as previously described^35^.

### EMSA

1. nM of Alexa647 labeled nucleic acids (IDT) were incubated with indicated protein concentrations in a total of 20 µl of EMSA binding buffer (20 mM HEPES pH 7.9, 150 mM NaCl, 1 mM MgCl2, 2.5

%, 0.5 mM TCEP, 1.3 ug/ul BSA) and incubated for 30 min on ice. 4 µl of EMSA loading buffer (50 % glycerol, 0.075 % bromophenol blue) was added to the samples before running them on a 2 % (w/v) agarose gel in 0.5 x TBE + 2.25 mM MgCl2 at 80 V for approximately 1.5 h. Gels were imaged on a ChemiDoc acquisition system (BioRad). Oligos used for EMSA assays are listed in Table S4.

### In-vitro droplet formation assay

PEGylated plates for microscopy were prepared as described^87^. Before performing droplet formation, wells were passivated using 100 mg/ml BSA for 30 min and rinsed with ddH2O. To induce condensate formation, proteins were prepared at 2x concentration in protein storage buffer (300 mM NaCl) and then diluted to 150 mM NaCl with ddH2O. For testing the effect of nucleic acids on condensate formation, the indicated concentrations were prepared as a 2x mix in ddH2O. Condensates were allowed to settle for 30min in PEGylated microcopy plates before imaging. Z-stacks consisting of 11 slices and a Z-stack step size of 0.64µm of 3 different areas per condition were acquired on an Olympus spinning disk confocal microscope equipped with a 20x/0.75 UPLSApo WD – 0.6 mm objective and Hamamatsu Orca Flash 4.0 camera. Focal plane for imaging was automatically determined using IX3 Z-Drift Compensator and the images were acquired using the 488nm laser. FRAP assays and RNA condensate colocalization was done on a Zeiss LSM-980 confocal microscope by imaging a single confocal plane.

### Quantification of condensates

Quantification of condensates was done in ImageJ. First a maximum intensity projection of Z-stack was made followed by background subtraction using a rolling ball radius of 50. Automatic thresholding was done using the Triangle algorithm and condensate area was quantified by analyzing particles with a minimum size of 2µm^2^ and a circularity of 0.9-1. Droplet area was plotted using R.

### Data and code availability statement

Coordinates and structure factors of LC8 in complex with Panx peptides harboring TQT site #1, site #2 or both have been deposited in the Protein Data Bank (PDB) under the accession number 7K3K, 7K3L and 7K3J, respectively. All sequencing data produced for this publication has been deposited to the NCBI GEO archive under the accession number GSE159424. The mass spectrometry proteomics data have been deposited to the ProteomeXchange Consortium via the PRIDE^88^ partner repository with the dataset identifier PXD022010.

---

**Figure S1:**
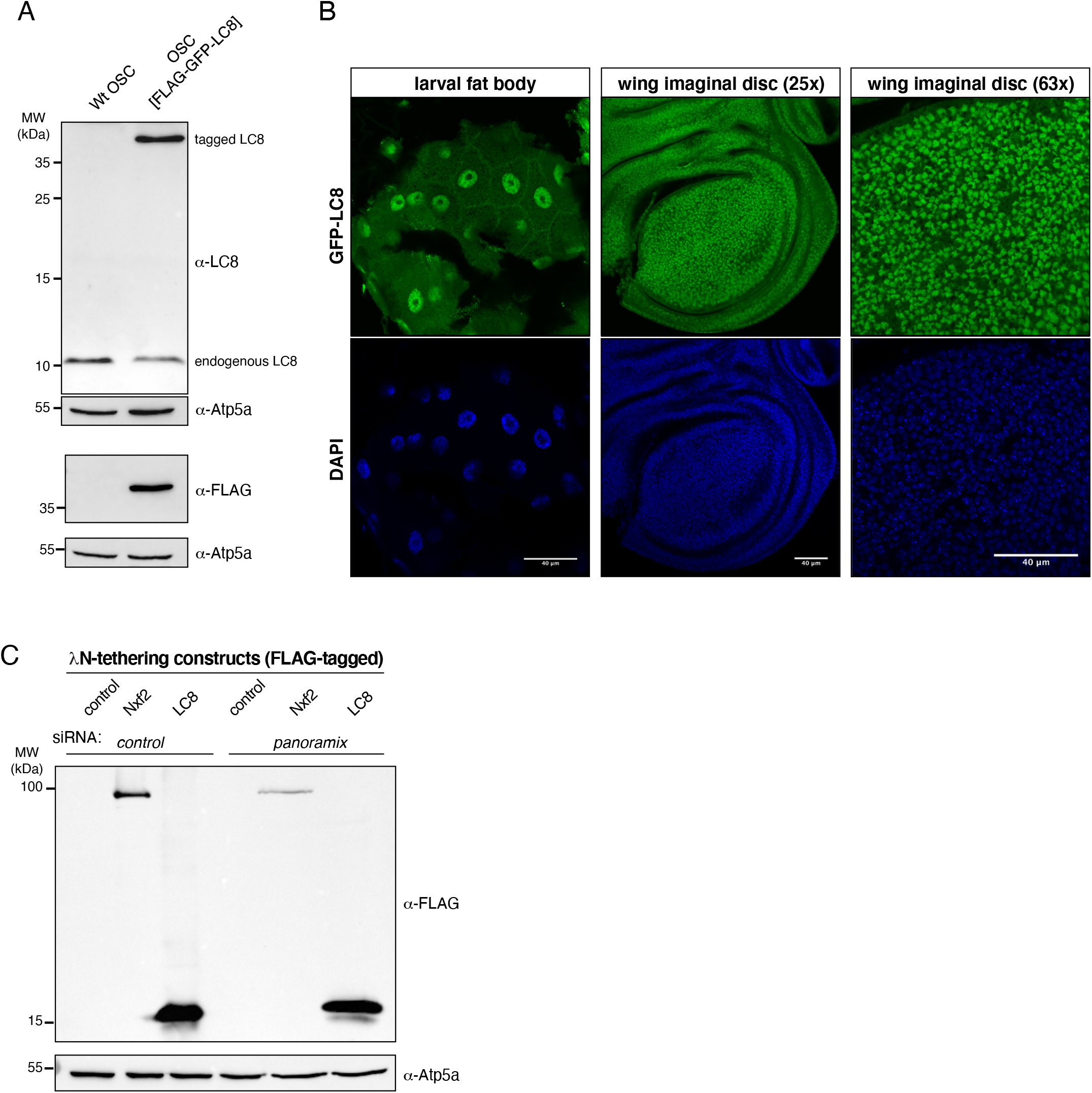
related to Figure 1. **A**, Western blot showing expression of endogenously FLAG-GFP tagged LC8 (one genomic copy) in a stable OSC line probed with anti-LC8 antibody (top) and anti-FLAG antibody (bottom; Atp5a served as loading control). **B**, Confocal images showing expression and nuclear localization of GFP-LC8 in different Drosophila larval tissues (blue DAPI staining indicates DNA; scale bars: 40µm). **C**, Western blot showing expression levels of indicated λN fusion proteins in transiently transfected OSCs which were co-transfected with control or panx siRNAs (related to Fig. 1F, Atp5a served as loading control).

**Figure S2:**
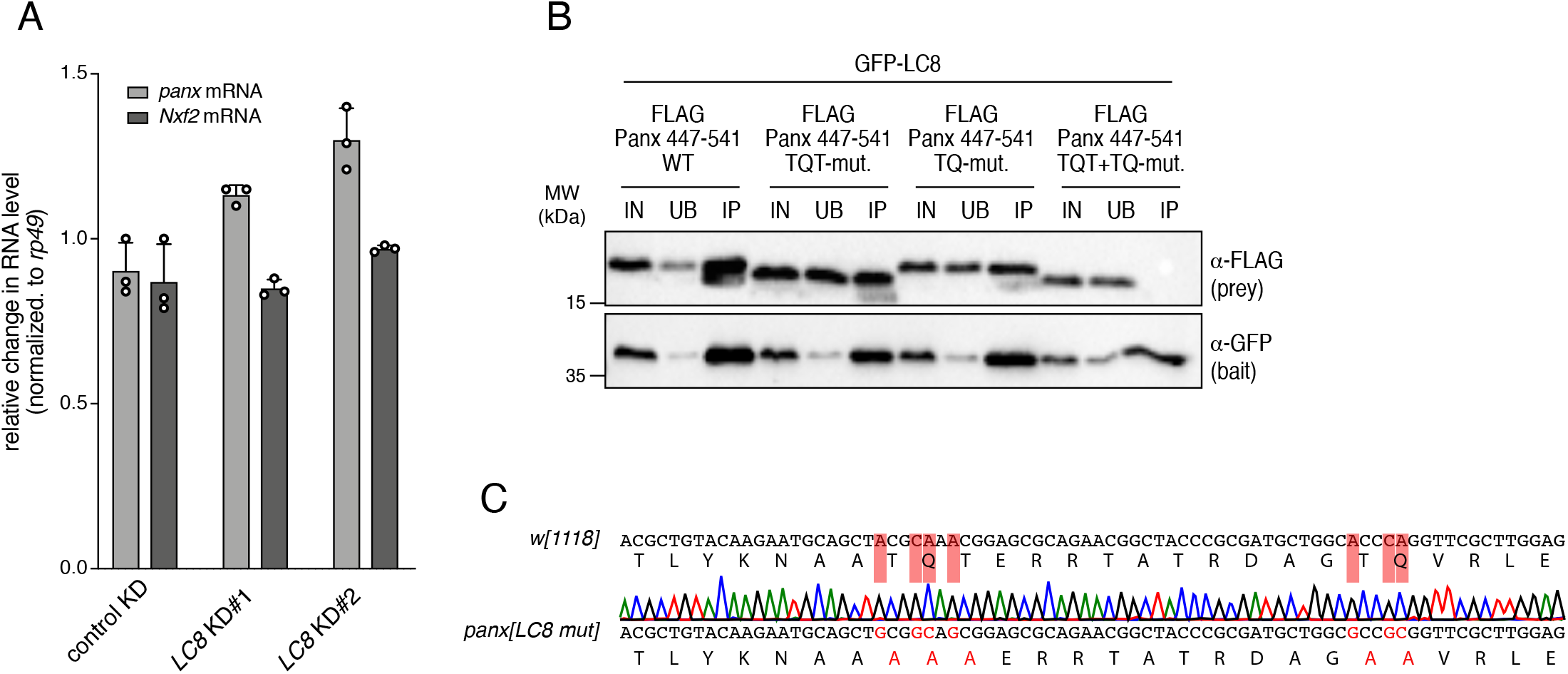
related to Figure 2. **A**, Bar graph showing mRNA levels of panx and nxf2 in OSCs after control depletion or depletion of LC8 (with 2 independent siRNAs; data represent mean + St.dev. of 3 independent experiments). **B**, Western blot analysis of GFP-LC8 immunoprecipitation experiments using lysate from S2 cells transiently transfected with indicated FLAG-Panoramix expression plasmids (relative amount loaded in immunoprecipitation lanes: 4x; IN: input, UB: unbound, IP; immono-precipitate). **C**, Sanger sequencing profile of genomic DNA and corresponding translation from panx[LC8-mut] flies (generated by CRISPR engineering). Mutations differing from the wildtype panx allele (w[1118] control flies) are highlighted in red.

**Figure S3:**
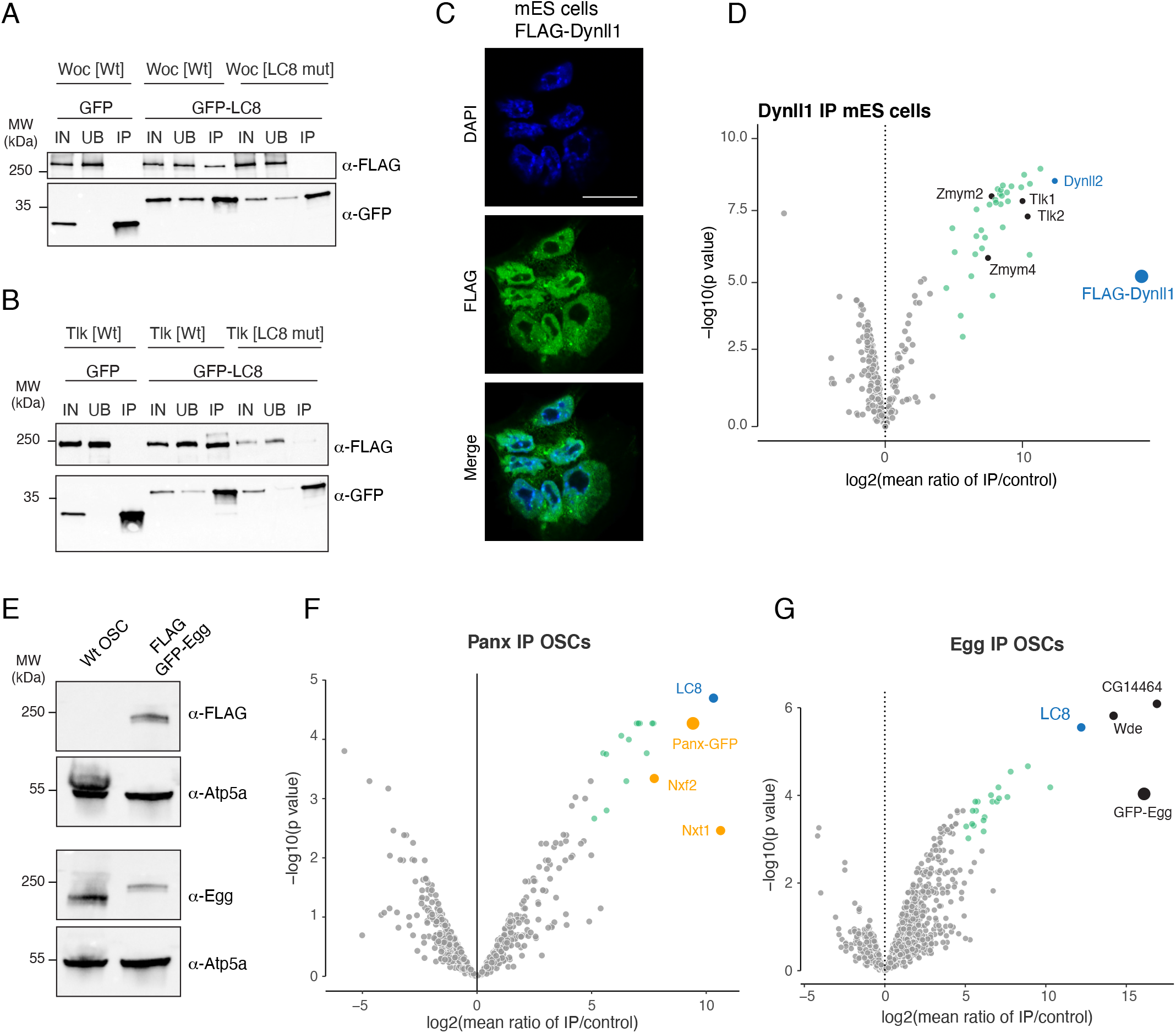
related to Figure 3. **A, B**, Western blot analyses of GFP or GFP-LC8 co-immunoprecipitation experiments using lysate from S2 cells transiently transfected with indicated FLAG-Woc (A) or FLAG-Tlk (B) expression plasmids (relative amount loaded in immunoprecipitation lanes: 4x; IN: input, UB: unbound, IP; immono-precipitate). **C**, Confocal images of mouse ES cells showing FLAG-Dynll1 (green) and DNA (blue; scale bar: 20µm). **D**, Volcano plot showing fold enrichments versus statistical significance (determined by quantitative mass spectrometry) of proteins in FLAG-Dynll1 co-immunoprecipitates versus control (n=3 biological replicates; experimental ES cells express endogenously tagged Dynll1; wildtype ES cells served as control). The bait (FLAG-Dynll1) as well as selected interacting nuclear proteins are labelled. **E**, Western blot indicating expression of endogenously FLAG-GFP tagged Eggless in an OSC line (probed with FLAG antibody (top) and anti-Eggless antibody (Atp5a served as loading control). **F, G**, Volcano plots showing fold enrichments versus statistical significance (determined by quantitative mass spectrometry) of proteins in FLAG-GFP-Panx co-immunoprecipitates (F) or in FLAG-GFP-Egg co-immunoprecipitates (G) versus control (n=3 biological replicates; experimental OSCs express endogenously tagged Panx (F) or Egg (G); wildtype OSCs served as control).

**Figure S4:**
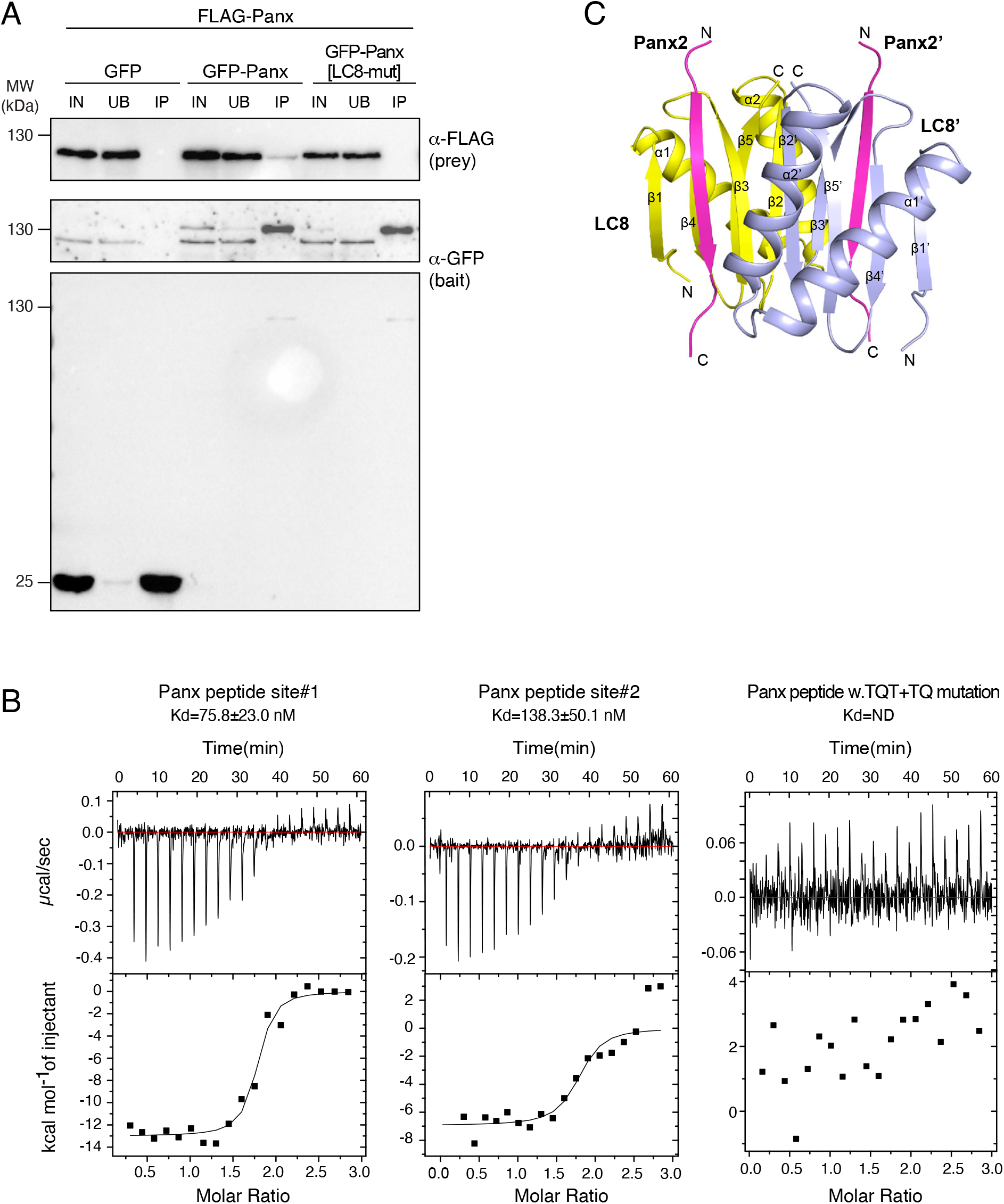
related to Figure 4. **A**, Western blot analysis of GFP or GFP-Panx immunoprecipitation experiments from OSCs lysates co-transfected with indicated Flag-tagged Panx variants. **B**, ITC binding curves for complex formation between LC8 homodimers and Panoramix peptides containing LC8 binding site #1 (left), site #2 (middle) or a peptide with bith sites harboring Alanne mutations in the TQT and TQ motifs. **C**, Ribbon representation of the structure of a LC8 homodimer (monomers in yellow and cyan) in complex with two Panoramix peptides containing the LC8 interacting site#1 (TQT site).

**Figure S5:**
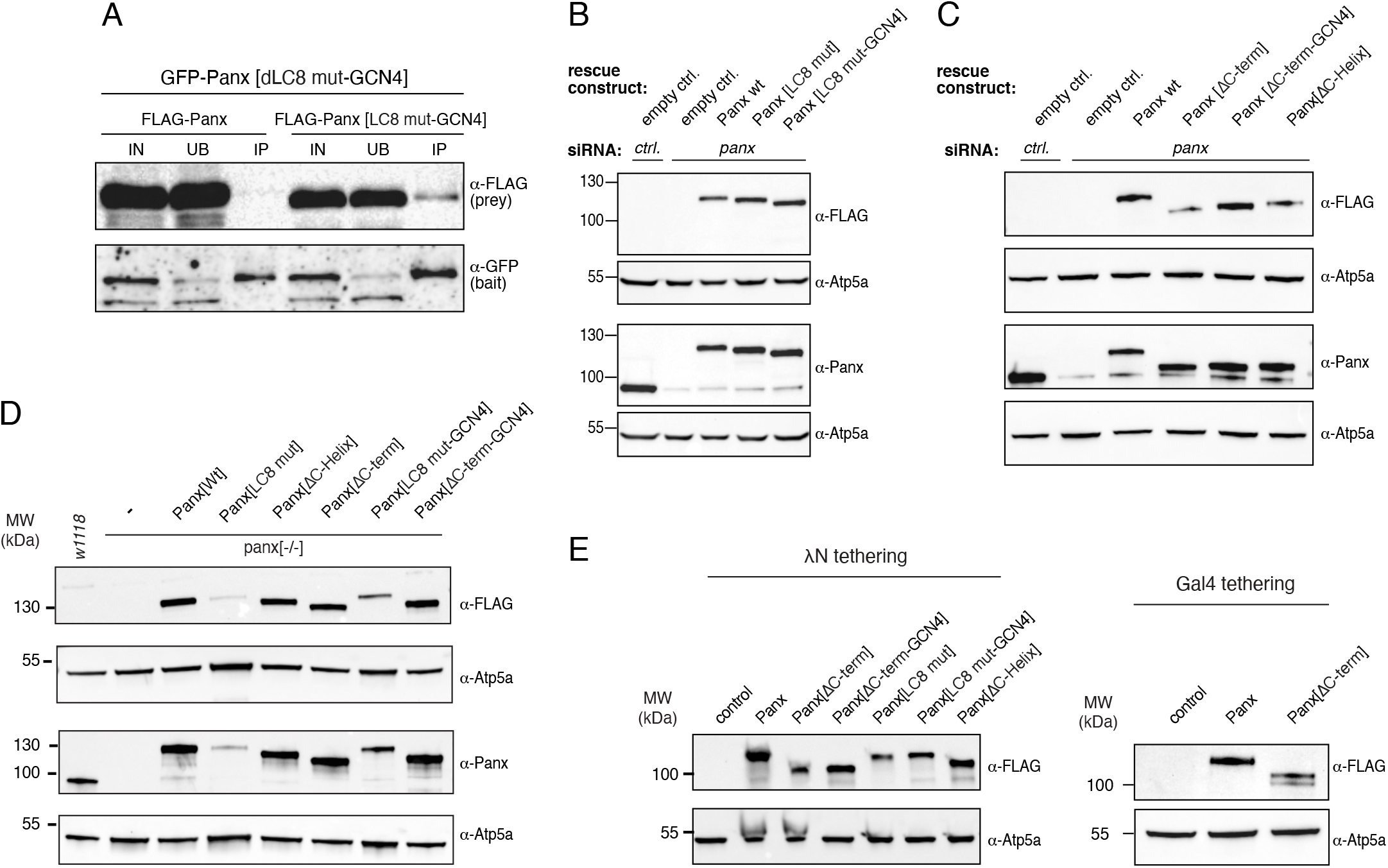
related to Figure 5. **A**, Western blot analysis of GFP-Panx[ΔLC8-GCN4] immunoprecipitation experiments from OSCs lysates cotransfected with indicated FLAG-tagged Panx variants.**B-C**, Western blots showing levels of indicated proteins in lysates from OSCs with indicated siRNA-mediated knockdowns and transiently transfected, siRNA-resistant rescue constructs (relates to experiment shown in Fig. 5A; Atp5A served as loading control). **D**, Western blots showing levels of indicated proteins in ovary lysates from flies with indicated genotypes (related to experiments shown in Fig. 5C-D; Atp5A served as loading control). **E**, Western blots showing levels of indicated λN-(left) and Gal4-(right) fusion proteins transiently expressed in OSCs carrying the GFP-reporter contruct (related to Fig. 5F, Atp5a served as loading control).

**Figure S6:**
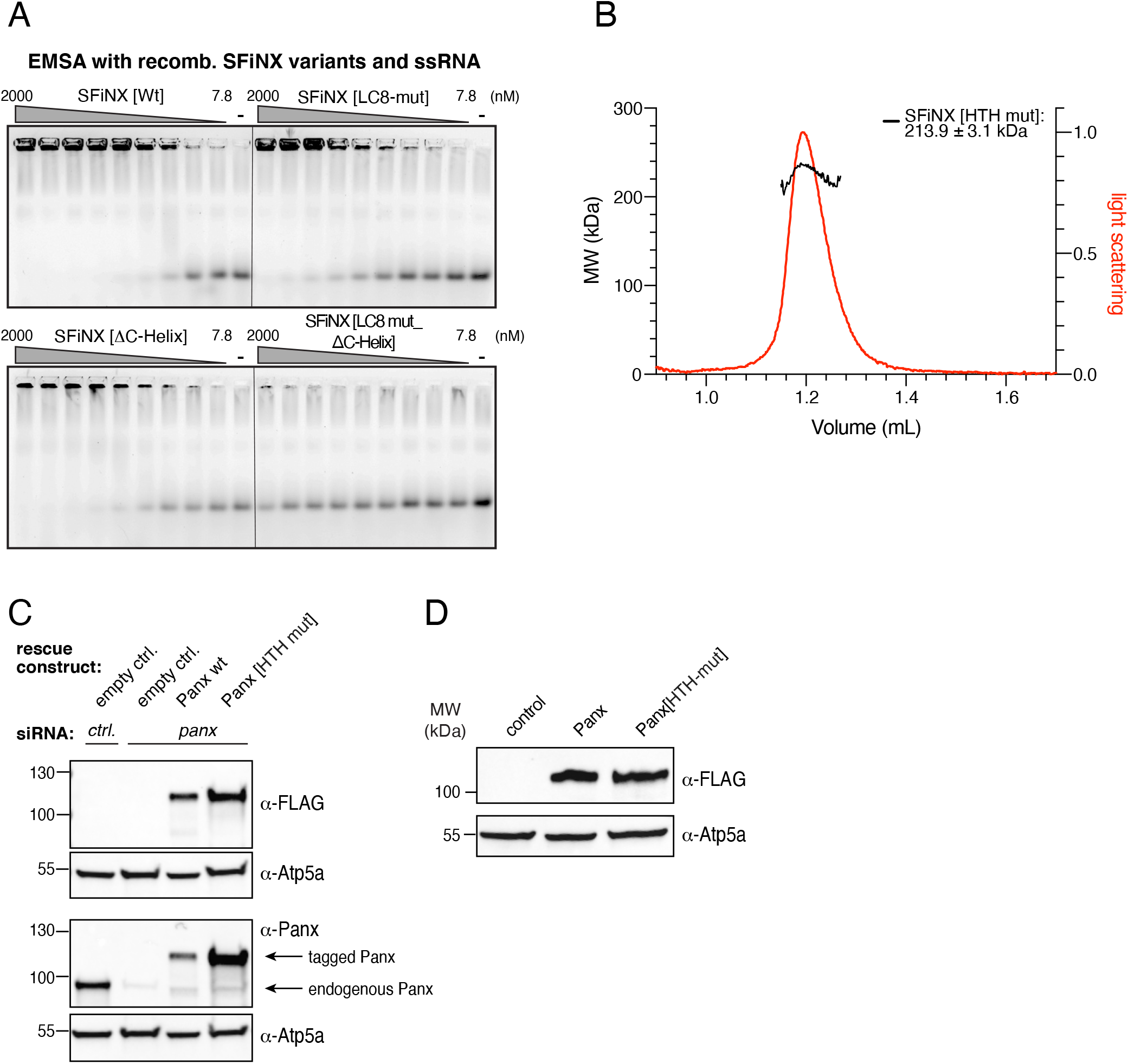
related to Figure 6. **A**, Agarose gel images showing Electro Mobility Shift Assays (EMSA) using single stranded RNA with indicated SFiNX complex variants. Serial dilution of recombinant complexes ranging from 7.8 to 2000 nM is shown (‘-’ indicates no protein). Quantification of data shown in Figure 6B. **B**, SEC-MALS chromatogram and determined molecular weight of recombinant SFiNX[HTH-mut] complex variants (related to Figure 6E). **C**, Western blots showing levels of indicated proteins in lysates from OSCs with indicated siRNA-mediated knockdowns and transiently transfected, siRNA-resistant rescue constructs (relates to experiment shown in Fig. 6E; Atp5A served as loading control). **D**, Western blots showing levels of indicated λN-fusion proteins transiently expressed in OSCs carrying the GFP-reporter contruct (related to Fig. 6F, Atp5a served as loading control).

**Figure S7:**
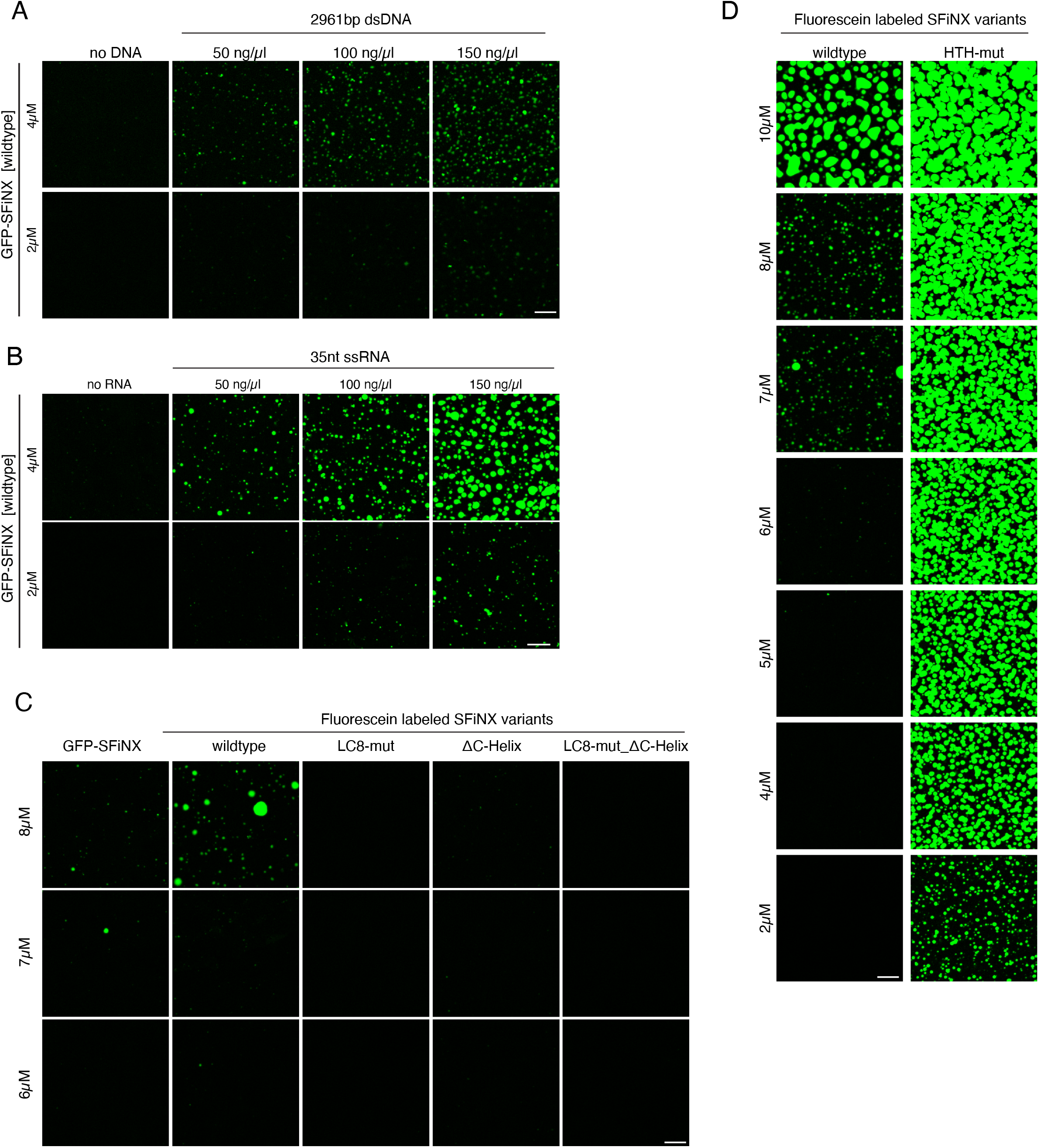
related to Figure 7. **A-B**, Confocal images showing enhanced condensate formation of mEGFP-SFiNX at indicated concentrations with increasing dsDNA (A) and ssRNA (B) concentration (scale bar: 20*µ*m). **C-D**, Confocal images showing in vitro condensate formation of indicated fluorescein-labeled recombinant SFiNX variants at indicated concentrations (scale bar: 20*µ*m).

## Notes

### Competing Interest Statement

The authors have declared no competing interest.

## References

1 Grewal, S. I. & Moazed, D. Heterochromatin and epigenetic control of gene expression. Science 301, 798–802, doi:10.1126/science.1086887 (2003).

2 Slotkin, R. K. & Martienssen, R. Transposable elements and the epigenetic regulation of the genome. Nature reviews. Genetics 8, 272–285 (2007).

3 Fedoroff, N. V. Presidential address. Transposable elements, epigenetics, and genome evolution. Science 338, 758–767 (2012).

4 Janssen, A., Colmenares, S. U. & Karpen, G. H. Heterochromatin: Guardian of the Genome. Annual review of cell and developmental biology 34, 265–288, doi:10.1146/annurev-cellbio-100617-062653 (2018).

5 Allshire, R. C. & Madhani, H. D. Ten principles of heterochromatin formation and function. Nature reviews. Molecular cell biology 19, 229–244, doi:10.1038/nrm.2017.119 (2018).

6 Rea, S. et al. Regulation of chromatin structure by site-specific histone H3 methyltransferases. Nature 406, 593–599, doi:10.1038/35020506 (2000).

7 Bannister, A. J. et al. Selective recognition of methylated lysine 9 on histone H3 by the HP1 chromo domain. Nature 410, 120–124, doi:10.1038/35065138 (2001).

8 Nakayama, J., Rice, J. C., Strahl, B. D., Allis, C. D. & Grewal, S. I. Role of histone H3 lysine 9 methylation in epigenetic control of heterochromatin assembly. Science 292, 110–113, doi:10.1126/science.1060118 (2001).

9 Lachner, M., O’Carroll, D., Rea, S., Mechtler, K. & Jenuwein, T. Methylation of histone H3 lysine 9 creates a binding site for HP1 proteins. Nature 410, 116–120, doi:10.1038/35065132 (2001).

10 Ninova, M., Fejes Toth, K. & Aravin, A. A. The control of gene expression and cell identity by H3K9 trimethylation. Development 146, doi:10.1242/dev.181180 (2019).

11 Yang, P., Wang, Y. & Macfarlan, T. S. The Role of KRAB-ZFPs in Transposable Element Repression and Mammalian Evolution. Trends Genet 33, 871–881, doi:10.1016/j.tig.2017.08.006 (2017).

12 Ghildiyal, M. & Zamore, P. D. Small silencing RNAs: an expanding universe. Nature reviews. Genetics 10, 94–108 (2009).

13 Holoch, D. & Moazed, D. RNA-mediated epigenetic regulation of gene expression. Nature reviews. Genetics 16, 71–84, doi:10.1038/nrg3863 (2015).

14 Grewal, S. I. RNAi-dependent formation of heterochromatin and its diverse functions. Current opinion in genetics & development 20, 134–141, doi:10.1016/j.gde.2010.02.003 (2010).

15 Martienssen, R. & Moazed, D. RNAi and heterochromatin assembly. Cold Spring Harb Perspect Biol 7, a019323, doi:10.1101/cshperspect.a019323 (2015).

16 Shimada, Y., Mohn, F. & Buhler, M. The RNA-induced transcriptional silencing complex targets chromatin exclusively via interacting with nascent transcripts. Genes & development 30, 2571–2580, doi:10.1101/gad.292599.116 (2016).

17 Siomi, M. C., Sato, K., Pezic, D. & Aravin, A. A. PIWI-interacting small RNAs: the vanguard of genome defence. Nature reviews. Molecular cell biology 12, 246–258, doi:nrm3089 [pii] 10.1038/nrm3089 (2011).

18 Ozata, D. M., Gainetdinov, I., Zoch, A., O’Carroll, D. & Zamore, P. D. PIWI-interacting RNAs: small RNAs with big functions. Nature reviews. Genetics, doi:10.1038/s41576-018-0073-3 (2018).

19 Czech, B. et al. piRNA-Guided Genome Defense: From Biogenesis to Silencing. Annual review of genetics 52, 131–157, doi:10.1146/annurev-genet-120417-031441 (2018).

20 Aravin, A. A., Hannon, G. J. & Brennecke, J. The Piwi-piRNA pathway provides an adaptive defense in the transposon arms race. Science 318, 761–764 (2007).

21 Senti, K. A. & Brennecke, J. The piRNA pathway: a fly’s perspective on the guardian of the genome. Trends Genet 26, 499–509, doi:S0168-9525(10)00171-X [pii] 10.1016/j.tig.2010.08.007 (2010).

22 Czech, B. & Hannon, G. J. One Loop to Rule Them All: The Ping-Pong Cycle and piRNA-Guided Silencing. Trends in biochemical sciences 41, 324–337, doi:10.1016/j.tibs.2015.12.008 (2016).

23 Cox, D. N., Chao, A. & Lin, H. piwi encodes a nucleoplasmic factor whose activity modulates the number and division rate of germline stem cells. Development 127, 503–514 (2000).

24 Saito, K. et al. Specific association of Piwi with rasiRNAs derived from retrotransposon and heterochromatic regions in the Drosophila genome. Genes & development 20, 2214–2222 (2006).

25 Brennecke, J. et al. Discrete Small RNA-Generating Loci as Master Regulators of Transposon Activity in Drosophila. Cell 128, 1089–1103 (2007).

26 Wang, S. H. & Elgin, S. C. Drosophila Piwi functions downstream of piRNA production mediating a chromatin-based transposon silencing mechanism in female germ line. Proceedings of the National Academy of Sciences of the United States of America 108, 21164–21169, doi:10.1073/pnas.1107892109 (2011).

27 Sienski, G., Donertas, D. & Brennecke, J. Transcriptional silencing of transposons by Piwi and maelstrom and its impact on chromatin state and gene expression. Cell 151, 964–980, doi:10.1016/j.cell.2012.10.040 (2012).

28 Le Thomas, A. et al. Piwi induces piRNA-guided transcriptional silencing and establishment of a repressive chromatin state. Genes & development 27, 390–399, doi:10.1101/gad.209841.112 (2013).

29 Rozhkov, N. V., Hammell, M. & Hannon, G. J. Multiple roles for Piwi in silencing Drosophila transposons. Genes & development 27, 400–412, doi:10.1101/gad.209767.112 (2013).

30 Yu, Y. et al. Panoramix enforces piRNA-dependent cotranscriptional silencing. Science 350, 339–342, doi:10.1126/science.aab0700 (2015).

31 Sienski, G. et al. Silencio/CG9754 connects the Piwi-piRNA complex to the cellular heterochromatin machinery. Genes & development 29, 2258–2271, doi:10.1101/gad.271908.115 (2015).

32 Iwasaki, Y. W. et al. Piwi Modulates Chromatin Accessibility by Regulating Multiple Factors Including Histone H1 to Repress Transposons. Molecular cell 63, 408–419, doi:10.1016/j.molcel.2016.06.008 (2016).

33 Ninova, M. et al. Su(var)2-10 and the SUMO Pathway Link piRNA-Guided Target Recognition to Chromatin Silencing. Molecular cell 77, 556–570 e556, doi:10.1016/j.molcel.2019.11.012 (2020).

34 Mugat, B. et al. The Mi-2 nucleosome remodeler and the Rpd3 histone deacetylase are involved in piRNA-guided heterochromatin formation. Nature communications 11, 2818, doi:10.1038/s41467-020-16635-5 (2020).

35 Batki, J. et al. The nascent RNA binding complex SFiNX licenses piRNA-guided heterochromatin formation. Nature structural & molecular biology 26, 720–731, doi:10.1038/s41594-019-0270-6 (2019).

36 Murano, K. et al. Nuclear RNA export factor variant initiates piRNA-guided co-transcriptional silencing. The EMBO journal 38, e102870, doi:10.15252/embj.2019102870 (2019).

37 Fabry, M. H. et al. piRNA-guided co-transcriptional silencing coopts nuclear export factors. Elife 8, doi:10.7554/eLife.47999 (2019).

38 Zhao, K. et al. A Pandas complex adapted for piRNA-guided transcriptional silencing and heterochromatin formation. Nature cell biology 21, 1261–1272, doi:10.1038/s41556-019-0396-0 (2019).

39 Niki, Y., Yamaguchi, T. & Mahowald, A. P. Establishment of stable cell lines of Drosophila germ-line stem cells. Proceedings of the National Academy of Sciences of the United States of America 103, 16325–16330 (2006).

40 Saito, K. et al. A regulatory circuit for piwi by the large Maf gene traffic jam in Drosophila. Nature 461, 1296–1299 (2009).

41 Phillis, R., Statton, D., Caruccio, P. & Murphey, R. K. Mutations in the 8 kDa dynein light chain gene disrupt sensory axon projections in the Drosophila imaginal CNS. Development 122, 2955–2963 (1996).

42 Dick, T., Ray, K., Salz, H. K. & Chia, W. Cytoplasmic dynein (ddlc1) mutations cause morphogenetic defects and apoptotic cell death in Drosophila melanogaster. Molecular and cellular biology 16, 1966–1977, doi:10.1128/mcb.16.5.1966 (1996).

43 Lo, K. W., Naisbitt, S., Fan, J. S., Sheng, M. & Zhang, M. The 8-kDa dynein light chain binds to its targets via a conserved (K/R)XTQT motif. The Journal of biological chemistry 276, 14059–14066, doi:10.1074/jbc.M010320200 (2001).

44 Makokha, M., Hare, M., Li, M., Hays, T. & Barbar, E. Interactions of cytoplasmic dynein light chains Tctex-1 and LC8 with the intermediate chain IC74. Biochemistry 41, 4302–4311, doi:10.1021/bi011970h (2002).

45 Jespersen, N. et al. Systematic identification of recognition motifs for the hub protein LC8. Life Sci Alliance 2, doi:10.26508/lsa.201900366 (2019).

46 Barbar, E. Dynein light chain LC8 is a dimerization hub essential in diverse protein networks. Biochemistry 47, 503–508, doi:10.1021/bi701995m (2008).

47 Rapali, P. et al. DYNLL/LC8: a light chain subunit of the dynein motor complex and beyond. FEBS J 278, 2980–2996, doi:10.1111/j.1742-4658.2011.08254.x (2011).

48 Erdos, G. et al. Novel linear motif filtering protocol reveals the role of the LC8 dynein light chain in the Hippo pathway. PLoS Comput Biol 13, e1005885, doi:10.1371/journal.pcbi.1005885 (2017).

49 Fan, J., Zhang, Q., Tochio, H., Li, M. & Zhang, M. Structural basis of diverse sequence-dependent target recognition by the 8 kDa dynein light chain. J Mol Biol 306, 97–108, doi:10.1006/jmbi.2000.4374 (2001).

50 Pfister, K. K. et al. Genetic analysis of the cytoplasmic dynein subunit families. PLoS genetics 2, e1, doi:10.1371/journal.pgen.0020001 (2006).

51 Thurmond, J. et al. FlyBase 2.0: the next generation. Nucleic acids research 47, D759–D765, doi:10.1093/nar/gky1003 (2019).

52 Font-Burgada, J., Rossell, D., Auer, H. & Azorin, F. Drosophila HP1c isoform interacts with the zinc-finger proteins WOC and Relative-of-WOC to regulate gene expression. Genes & development 22, 3007–3023, doi:10.1101/gad.481408 (2008).

53 Kessler, R. et al. dDsk2 regulates H2Bub1 and RNA polymerase II pausing at dHP1c complex target genes. Nature communications 6, 7049, doi:10.1038/ncomms8049 (2015).

54 Sillje, H. H. & Nigg, E. A. Identification of human Asf1 chromatin assembly factors as substrates of Tousled-like kinases. Current biology: CB 11, 1068–1073, doi:10.1016/s0960-9822(01)00298-6 (2001).

55 Osumi, K., Sato, K., Murano, K., Siomi, H. & Siomi, M. C. Essential roles of Windei and nuclear monoubiquitination of Eggless/SETDB1 in transposon silencing. EMBO reports 20, e48296, doi:10.15252/embr.201948296 (2019).

56 Koch, C. M., Honemann-Capito, M., Egger-Adam, D. & Wodarz, A. Windei, the Drosophila homolog of mAM/MCAF1, is an essential cofactor of the H3K9 methyl transferase dSETDB1/Eggless in germ line development. PLoS genetics 5, e1000644, doi:10.1371/journal.pgen.1000644 (2009).

57 Mutlu, B. et al. Regulated nuclear accumulation of a histone methyltransferase times the onset of heterochromatin formation in C. elegans embryos. Sci Adv 4, eaat6224, doi:10.1126/sciadv.aat6224 (2018).

58 Kidane, A. I. et al. Structural features of LC8-induced self-association of swallow. Biochemistry 52, 6011–6020, doi:10.1021/bi400642u (2013).

59 Liang, J., Jaffrey, S. R., Guo, W., Snyder, S. H. & Clardy, J. Structure of the PIN/LC8 dimer with a bound peptide. Nat Struct Biol 6, 735–740, doi:10.1038/11501 (1999).

60 Gallego, P., Velazquez-Campoy, A., Regue, L., Roig, J. & Reverter, D. Structural analysis of the regulation of the DYNLL/LC8 binding to Nek9 by phosphorylation. The Journal of biological chemistry 288, 12283–12294, doi:10.1074/jbc.M113.459149 (2013).

61 Wang, L., Hare, M., Hays, T. S. & Barbar, E. Dynein light chain LC8 promotes assembly of the coiled-coil domain of swallow protein. Biochemistry 43, 4611–4620, doi:10.1021/bi036328x (2004).

62 O’Shea, E. K., Klemm, J. D., Kim, P. S. & Alber, T. X-ray structure of the GCN4 leucine zipper, a two-stranded, parallel coiled coil. Science 254, 539–544, doi:10.1126/science.1948029 (1991).

63 Goldman, C. H., Neiswender, H., Veeranan-Karmegam, R. & Gonsalvez, G. B. The Egalitarian binding partners Dynein light chain and Bicaudal-D act sequentially to link mRNA to the Dynein motor. Development 146, doi:10.1242/dev.176529 (2019).

64 Alberti, S., Gladfelter, A. & Mittag, T. Considerations and Challenges in Studying Liquid-Liquid Phase Separation and Biomolecular Condensates. Cell 176, 419–434, doi:10.1016/j.cell.2018.12.035 (2019).

65 Erdel, F. et al. Mouse Heterochromatin Adopts Digital Compaction States without Showing Hallmarks of HP1-Driven Liquid-Liquid Phase Separation. Molecular cell 78, 236–249 e237, doi:10.1016/j.molcel.2020.02.005 (2020).

66 Post, C., Clark, J. P., Sytnikova, Y. A., Chirn, G. W. & Lau, N. C. The capacity of target silencing by Drosophila PIWI and piRNAs. Rna 20, 1977–1986, doi:10.1261/rna.046300.114 (2014).

67 Verdel, A. et al. RNAi-mediated targeting of heterochromatin by the RITS complex. Science 303, 672–676 (2004).

68 Maksimov, V. et al. The binding of Chp2’s chromodomain to methylated H3K9 is essential for Chp2’s role in heterochromatin assembly in fission yeast. PloS one 13, e0201101, doi:10.1371/journal.pone.0201101 (2018).

69 Li, H. et al. An alpha motif at Tas3 C terminus mediates RITS cis spreading and promotes heterochromatic gene silencing. Molecular cell 34, 155–167, doi:10.1016/j.molcel.2009.02.032 (2009).

70 Cerase, A. et al. Phase separation drives X-chromosome inactivation: a hypothesis. Nature structural & molecular biology 26, 331–334, doi:10.1038/s41594-019-0223-0 (2019).

71 Pandya-Jones, A. et al. A protein assembly mediates Xist localization and gene silencing. Nature, doi:10.1038/s41586-020-2703-0 (2020).

72 Daneshvar, K. et al. lncRNA DIGIT and BRD3 protein form phase-separated condensates to regulate endoderm differentiation. Nature cell biology, doi:10.1038/s41556-020-0572-2 (2020).

73 Rapali, P. et al. Directed evolution reveals the binding motif preference of the LC8/DYNLL hub protein and predicts large numbers of novel binders in the human proteome. PloS one 6, e18818, doi:10.1371/journal.pone.0018818 (2011).

74 Venken, K. J. et al. Versatile P[acman] BAC libraries for transgenesis studies in Drosophila melanogaster. Nature methods 6, 431–434, doi:10.1038/nmeth.1331 (2009).

75 Ejsmont, R. K., Bogdanzaliewa, M., Lipinski, K. A. & Tomancak, P. Production of fosmid genomic libraries optimized for liquid culture recombineering and cross-species transgenesis. Methods in molecular biology 772, 423–443, doi:10.1007/978-1-61779-228-1_25 (2011).

76 Gokcezade, J., Sienski, G. & Duchek, P. Efficient CRISPR/Cas9 Plasmids for Rapid and Versatile Genome Editing in Drosophila. G3, doi:10.1534/g3.114.014126 (2014).

77 Pfeiffer, B. D. et al. Tools for neuroanatomy and neurogenetics in Drosophila. Proceedings of the National Academy of Sciences of the United States of America 105, 9715–9720, doi:10.1073/pnas.0803697105 (2008).

78 Meers, M. P., Bryson, T. D., Henikoff, J. G. & Henikoff, S. Improved CUT&RUN chromatin profiling tools. Elife 8, doi:10.7554/eLife.46314 (2019).

79 Langmead, B. & Salzberg, S. L. Fast gapped-read alignment with Bowtie 2. Nature methods 9, 357–359, doi:10.1038/nmeth.1923 (2012).

80 Shen, L., Shao, N., Liu, X. & Nestler, E. ngs.plot: Quick mining and visualization of next-generation sequencing data by integrating genomic databases. BMC Genomics 15, 284, doi:10.1186/1471-2164-15-284 (2014).

81 Elling, U. et al. A reversible haploid mouse embryonic stem cell biobank resource for functional genomics. Nature 550, 114–118, doi:10.1038/nature24027 (2017).

82 Subramanian, A. et al. Gene set enrichment analysis: a knowledge-based approach for interpreting genome-wide expression profiles. Proceedings of the National Academy of Sciences of the United States of America 102, 15545–15550, doi:10.1073/pnas.0506580102 (2005).

83 Battye, T. G., Kontogiannis, L., Johnson, O., Powell, H. R. & Leslie, A. G. iMOSFLM: a new graphical interface for diffraction-image processing with MOSFLM. Acta crystallographica. Section D, Biological crystallography 67, 271–281, doi:10.1107/S0907444910048675 (2011).

84 Kabsch, W. Integration, scaling, space-group assignment and post-refinement. Acta crystallographica. Section D, Biological crystallography 66, 133–144, doi:10.1107/S0907444909047374 (2010).

85 Adams, P. D. et al. PHENIX: building new software for automated crystallographic structure determination. Acta crystallographica. Section D, Biological crystallography 58, 1948–1954 (2002).

86 Emsley, P., Lohkamp, B., Scott, W. G. & Cowtan, K. Features and development of Coot. Acta crystallographica. Section D, Biological crystallography 66, 486–501, doi:10.1107/S0907444910007493 (2010).

87 Gibson, B. A. et al. Organization of Chromatin by Intrinsic and Regulated Phase Separation. Cell 179, 470–484 e421, doi:10.1016/j.cell.2019.08.037 (2019).

88 Vizcaino, J. A. et al. 2016 update of the PRIDE database and its related tools. Nucleic acids research 44, D447–456, doi:10.1093/nar/gkv1145 (2016).

